# Covert RNA viruses in medflies differ in their mode of transmission and tissue tropism

**DOI:** 10.1101/2023.11.08.566245

**Authors:** Luis Hernández-Pelegrín, Hannah-Isadora Huditz, Pablo García-Castillo, Norbert C.A. de Ruijter, Monique M. van Oers, Salvador Herrero, Vera I.D. Ros

**Author notes:** Correspondence to: <u>Salvador Herrero</u> Universitat de València. Department of Genetics Dr Moliner 50, 46100 Burjassot, Spain, <u>Vera I.D. Ros</u> Laboratory of Virology, Wageningen University and Research Droevendaalsesteeg 1, 6708 PB, Wageningen, The Netherlands.

## Abstract

Numerous studies have demonstrated the presence of covert viral infections in insects. These infections can be transmitted in insect populations via two main routes: vertical from parents to offspring, or horizontal between nonrelated individuals. Thirteen covert RNA viruses have been described in the Mediterranean fruit fly (medfly). Some of these viruses are established in different laboratory-reared and wild medfly populations, although variations in the viral repertoire and viral levels have been observed at different time points. To better understand these viral dynamics, we characterized the prevalence and levels of covert RNA viruses in two medfly strains, assessed the route of transmission of these viruses, and explored their distribution in medfly adult tissues. Altogether, our results indicated that the different RNA viruses found in medflies vary in their preferred route of transmission. Two iflaviruses and a narnavirus are predominantly transmitted through vertical transmission via the female, while a nodavirus and a nora virus exhibited a preference for horizontal transmission. Overall, our results give valuable insights into the viral tropism and transmission of RNA viruses in the medfly, contributing to the understanding of viral dynamics in insect populations.

## Background

As the most diversified group across the animal kingdom, insects are considered a reservoir of a great variety of viruses (Shi et al., 2016a). In recent times, the advent of high-throughput sequencing methods has led to the discovery of numerous RNA viruses, which has redefined the virome of insects (Bonning, 2020; Haoming et al., 2021; Käfer et al., 2019; Porter et al., 2019; Shi et al., 2016b). Most of these RNA viruses are insect-specific viruses (ISVs), with insects as their unique host, and cause covert viral infections, which are characterized by the absence of noticeable or lethal symptoms. However, covert infections can affect certain host fitness traits (Longdon et al., 2012; Williams et al., 2017; Yuan et al., 2020) or induce behavioral and physiological changes in the insect host (Fujiyuki et al., 2005; Han et al., 2015).

Regardless of the effects on the host associated with covert infections, intriguing questions are how covert infections are maintained for generations in insect populations and how they are transmitted. Viral transmission can occur from parents to offspring (vertically), between (not necessarily related) individuals often of the same species (horizontally); or a combination of both as reviewed by Shapiro-Ilan et al., 2012. Specifically, vertical transmission has been proposed to be a key factor for maintaining covert viruses in nature (Agboli et al., 2019), although few studies have provided direct experimental evidence. Vertical transmission can occur via sperm (paternal) or via the egg (maternal), and maternal transmission is subdivided into transovarial transmission, in which the pathogen is transmitted within the egg, or transovum transmission, in which the pathogen remains on the surface of the egg and infects the offspring after hatching (Solter & Becnel, 2017).

Up to date, thirteen viruses belonging to different families of positive single stranded (ss)RNA, negative ssRNA and double stranded (ds)RNA viruses have been discovered in the agricultural pest *Ceratitis capitata* (Wiedemann) (Diptera: Tephritidae), also known as the Mediterranean fruit fly or medfly (Hernández-Pelegrín et al., 2022; Kondo et al., 2019; Llopis-Giménez et al., 2017; Sharpe et al., 2021). These viruses co-infect in different combinations field-derived, laboratory-reared, and mass-reared medfly strains causing covert infections (Hernández-Pelegrín et al., 2022). Among medfly RNA viruses, Ceratitis capitata nora virus (CcaNV) is the best described virus. CcaNV infection negatively influences medfly fitness in terms of a reduced pupal weight, shorter adult survival under stress, and higher susceptibility to parasitism by *Aganaspis daci* [Hymenoptera: Figitidae] (Hernández-Pelegrín et al., 2023). CcaNV can be horizontally transmitted after the addition of purified virus into the larval diet (Llópis-Giménez et al., 2017; Hernández-Pelegrín et al., 2023). Yet besides CcaNV, an in-depth analysis of the distribution and transmission of covert RNA viruses infecting medfly is lacking.

From a practical point of view, the insect mass-rearing industry would benefit from a better understanding of insect virus ecology, since it is highly dependent on the optimal health of the insects. In this industry, the presence of pathogens represents a huge risk, especially in the artificial environmental conditions common in mass-rearing facilities (Eilenberg et al., 2015; Slowik et al., 2023). Specifically, covert RNA viruses can be easily transmitted between the conspecific reared insects and, in some scenarios, cause disease outbreaks (Bertola & Mutinelli, 2021; Maciel-Vergara & Ros, 2017). In the case of the medfly, the systematic area-wide release of sterile males produced in mass-rearing facilities is required for the application of the sterile insect technique (SIT), which is the most used strategy for the biocontrol of this horticultural pest (Enkerlin, 2005). Moreover, the presence of covert RNA viruses may alter the insect population dynamics in the field and in both cases, mass-rearing and field conditions, viral transmission is assumed to play a fundamental role.

In this article, we combined multiple approaches for the study of viral tissue tropism and transmission routes with the aim of gaining insights into the dynamic of insect-virus interactions, and determining if specific transmission mechanisms are related with the establishment and maintenance of covert infection with RNA viruses.

## Methods

### Insects

Two medfly strains, Vienna 8A (V8A) and Madrid, were used in the experiments. The V8A strain is produced for the application of sterile insect technique programs by the state-owned company Empresa de Transformacion Agraria S.A (Grupo TRAGSA, Valencia, Spain) at the mass-rearing facility in Caudete de las Fuentes (Valencia, Spain). The V8A strain, which is the result of a mix between the Vienna-8 mix 2002 strain and wild individuals collected in citrus orchards located in the province of Valencia (Spain), is maintained at 25 ± 1°C, 65% humidity, and 14/10 h light/dark cycles (Plá et al., 2021; Porras et al., 2020). The Madrid strain was established in 2001 with wild flies collected from experimental fields located at the Instituto Valenciano de Investigaciones Agrarias (IVIA). Since then, the colony has been reared under laboratory conditions of 26 °C, 40-60 % humidity, and 14/10 h light/dark cycles (Arouri et al., 2015). For both medfly strains, larval diet is composed of wheat bran, sugar, and brewe’s yeast (de Pedro et al., 2021) while adults are provided with water and a sugar-based sources *ad libitum*.

### Viral RNA levels in different medfly developmental stages

Larvae, pupae, and adults deriving from a single generation of both Madrid and V8A strains were obtained. Larvae were collected as third instar, pupae between 5 and 8 days after pupation and adults between 5-6 days after emergence. Six samples containing five individuals per sample were tested per developmental stage. For the adults, three samples contained only males, and three contained only females. RNA levels for the 13 RNA viruses previously described in medfly were determined for each sample using reverse transcription quantitative PCR (RT-qPCR) and normalized relatively to host gene expression (described below).

### Viral prevalence and levels in individual medfly adults

Thirty pupae per strain were individually placed in 2 ml microcentrifuge tubes. Per strain, 18 adults (nine males and nine females) were collected after emergence to determine the levels of RNA viruses. Normalized viral RNA levels of the five viruses known to be present in Madrid and V8A medfly strains were determined for each sample using RT-qPCR as described below.

A symmetric Venn Diagram was generated to visualize the repertoire of RNA viruses co-infecting single medfly adults (https://bioinformatics.psb.ugent.be/webtools/Venn/, accessed in April 2023). Moreover, we compared the observed and expected proportions of each possible viral pair to determine whether some viral combinations were favored or unfavored in the flies. Observed proportions were defined as the number of individuals presenting a given pair of viruses divided by the total number of individuals. Expected proportions for a given pair of viruses were calculated by multiplying the proportions observed for each virus individually. The statistical differences between observed and expected proportions were investigated using a Fishe’s exact test.

### Viral RNA levels in sterilized and non-sterilized medfly eggs

To explore the vertical transmission of RNA viruses, we assessed the viral prevalence and viral RNA levels in two groups of eggs: non-sterilized and sterilized. The eggs selected for the analysis were oviposited by the siblings of those medfly adults employed for the analysis of viral prevalence and viral RNA levels (see above). For surface sterilization, 100 µl of eggs were submerged into a 5% sodium hypochlorite-milliQ water solution for three minutes to generate the sterilized group. Additionally, 100 µl eggs were submerged in milliQ water for three minutes to generate the non-sterilized group. Then, eggs were washed during two minutes in a milliQ water solution. The washing step was repeated twice. Finally, medfly eggs were washed using PBS during one minute before RNA isolation. Viral levels of the five RNA viruses previously detected were investigated in three replicates of sterilized and non-sterilized eggs from the Madrid and the V8A strains using RT-qPCR as described below.

### RNA isolation and quantification of RNA viruses in the medfly

The levels of RNA viruses were determined using molecular methods as previously described in Hernández-Pelegrín et al. (2022). Briefly, total RNA was extracted using TriPure isolation reagent (cat. No. 11667157001; Roche) following the manufacture’s protocol. The isolated RNA was DNAse-treated (cat. No. 10694233; Invitrogen^TM^) and reversed transcribed into cDNA using oligo (dT) primers and random hexamers (Prime-Script RT Reagent Kit, Takara Bio Inc.). Then, the levels of the 13 covert RNA viruses previously described in the medfly, as well as the expression of the medfly gene encoding the ribosomal protein L23a (Genbank acc: XM004518966), were assessed through RT-qPCR (CFX96 Touch Real-Time PCR Detection System, BioRad) using specific primers (Hernández-Pelegrín et al., 2022; Llopis-Giménez et al., 2017) (Table S1). The normalized viral RNA levels refer to the levels of viral molecules in relation to the levels of the reference gene *L23a*, and were calculated by comparing Ct values of RNA viruses and L23a, after adjusting for primer efficiency (Herrero et al., 2019).

### Visualization and statistical analyses

Normalized viral RNA levels were analyzed and visualized using GraphPad Prism version 8.0.0 for Windows (GraphPad Software, San Diego, California USA, www.graphpad.com). For the statistical analysis, a normalized viral RNA level below to 10^-6^ viral molecules per L23a molecules was assigned for non-detected viruses, and values of normalized viral RNA levels were transformed to a logarithmic scale to avoid misinterpreting the differences between samples with normalized viral RNA levels with a value smaller than 1. The normality of the data was investigated using a Shapiro-Wilk test. Then, a two-way ANOVA test or a non-parametric Kruskal-Wallis test was applied to determine the differences in viral RNA levels between developmental stages or adult tissues, respectively. Pairwise comparisons between the means of each group (i.e. larval and pupal stage or gut and brain tissues) were performed using Tukeýs post-hoc tests (ANOVA) or Dunńs (Kruskal-Wallis) post-hoc tests (Table S3). An unpaired t-test was applied to compare sterilized and non-sterilized eggs employed for the analysis of viral vertical transmission via the eggs. Differences between samples were considered significant for p-values lower than 0.05.

### Viral transmission after co-habitation and mating of Madrid and V8A adults

To study whether the RNA viruses are vertically transmitted from parents to offspring, and whether they are horizontally transmitted between parents, thirty pairs of virgin male and female adults were established. Half of the mating pairs contained a male of the Madrid strain with a female of the V8A strain, and the other half contained a female of the Madrid strain with a male of the V8A strain. Mating pairs were placed into rearing cups modified with a net on the lid to facilitate oviposition, and with sugar and water provided *ad libitum*. After nine days of cohabitation, the adults were removed and frozen individually at – 20 °C prior to RNA isolation. The eggs derived from each mating pair were collected at two consecutive days and placed onto artificial diet, in which the hatching larvae shared food resources during development. Twenty-two mating pairs successfully produced offspring (eleven for Madrid male x V8A female and eleven for V8A male x Madrid female). Viral RNA levels were assessed in both adults of the successful mating pairs using RT-qPCR as described above. Ten mating pairs (five per combination) were selected (those with the largest difference in viral RNA levels between the male and the female) to determine the viral RNA levels in six pupae of the offspring.

### Detection of the viral replicative RNA strand by RT-qPCR

To assess whether after potential horizontal transmission during mating the virus is replicating in the receptive host, we tested the presence of the viral negative strand (intermediary molecule during the viral replication) by RT-qPCR. To that aim, for each virus screened we selected four of the mating pairs. After RNA isolation, 500 ng of total RNA was treated with DNAse (cat. No. 10694233; Invitrogen^TM^) and submitted to cDNA synthesis with the Prime Script RT Reagent Kit (Takara Bio Inc.) at 42°C for 30min. For the cDNA synthesis, specific primers complementary to the negative strand of each RNA virus and containing a 5’overhang to avoid self-priming (Table S1) were used. Negative controls lacking the RT enzyme or lacking the primers with the 5’overhang were added to confirm the specificity of the viral negative strand amplification (Figure S1). Three microliter of the cDNA was used for the PCR reaction with DreamTaq DNA polymerase (Thermo Fisher Scientific), using a forward primer annealing to the overhang of the primer used to make the cDNA and a reverse primer specific for each virus (Table S1). The PCR reaction was performed as follows: one step of 95 °C for five minutes, 35 cycles of annealing at 52 °C for 30 s, and elongation at 72 °C for one minute (Herrero et al., 2019). PCR results were visualized in 1% agarose gels. Two PCR negative controls were added, one lacking medfly cDNA and one containing medfly cDNA previously tested to be virus-free.

### Tissue tropism in adult medflies

Adults of both sexes were collected from the Madrid and V8A rearing cages on days 5 and 6 post emergence. Flies were chilled on ice, fixed on a wax pan using entomological pins and dissected under the binocular with the help of entomological tweezers (Dumont nr. 5). The dissected tissues were collected in RNA*later* solution (Sigma Aldrich R0901, St. Louis, MO, USA) to conserve the integrity of the RNA until RNA isolation. Four somatic tissues were collected per fly: brain, gut, crop, and legs. In addition, ovaries were collected from females and testes from males. For each tissue type, we analysed the virus levels in six samples each consisting of six pooled tissues as described above. For somatic tissues, half of the samples were collected from male adults and the other half from female adults.

### In situ detection of CcaIV2 and CcaNV in adult tissues using fluorescence in situ hybridization

Custom probe sets annealing to the sequences of Ceratitis capitata iflavirus 2 (CcaIV2) or Ceratitis capitata nora virus (CcaNV) were designed using the Stellaris® FISH probe designer (www.biosearchtech.com/support/tools/design-software/stellaris-probe-designer, accessed on 02 November 2022) and following the parameters described by Meki et al. (2021). A total of 48 probes, with a length between 19-21 nt each, were designed for each virus (Figure S2) (Orjalo et al., 2011). For visualization, Quasar 570 (Q570) dye was bound at the 5’ end of the CcaNV probes and Quasar 670 (Q670) to the 5’end of the CcaIV2 probes.

Adult medflies from the V8A strain were collected 5 to 6 days after emergence, chilled on ice and dissected in PBS to obtain brain, legs, gut, crop, ovaries, and testes as described above. Dissected tissues were immediately submerged in 3.7% paraformaldehyde (PFA) solution and maintained on ice until the end of all dissections. Then the tissues were whole mount incubated in the 3.7% PFA solution under constant shaking at room temperature (RT) for 20 minutes. After fixation, the tissues were washed once in PBS for ten minutes, and twice in PBS-T (1 x PBS with 0.3% TritonX-100) for ten minutes each. Then, tissues were rinsed once with 70% ethanol and incubated overnight in fresh 70% ethanol at 4°C with constant shaking. The following day the ethanol was substituted with sterile PBS, after which the PBS was replaced with pre-heated Stellaris® RNA FISH Wash Buffer A (cat no. SMF-WA1-60; LGC Biosearch Technologies), wherein the tissues were incubated for five minutes at RT to allow rehydration. For the hybridization step, the Stellaris® RNA FISH Wash Buffer A was substituted for 100 µL of the hybridization solution and the tubes were incubated in a humid dark chamber, overnight at 37°C. The hybridization solution contained, per mL, 890 µL of Stellaris® RNA FISH Hybridization Buffer (cat no. SMF-HB1-10; LGC Biosearch Technologies), 100 µL of deionized formamide and 10 µL of selected custom probe sets. Three hybridization solutions were prepared for the assay: one containing probes for CcaIV2 detection, one with probes for CcaNV detection, and one containing both sets of probes. After hybridization, samples were briefly washed three times with pre-heated Stellaris® RNA FISH Wash Buffer A and incubated for 30 minutes in the humid dark chamber at 37°C. To perform the F-actin staining, the Stellaris® RNA FISH Wash Buffer A was replaced by 100 µL of PBS containing Alexa Fluor ® 488 phalloidin (Invitrogen, Paisley, UK) in the ratio 100:1 and the samples were incubated at RT in the dark under constant shaking. Finally, the PBS solution containing Alexa Fluor ® 488 phalloidin was substituted by clean PBS, and the tissues were mounted in a drop of Roti-Mount© FluorCare, containing DAPI (4′,6-diamidino-2-phenylindole) (Carl Roth, Germany) for DNA staining on the RNA-FISH labelled slides.

A Leica Stellaris 5 Confocal LSM with Las X software V4.40 (Leica Microsystems) was used to localize and spectrally separate the labels. By using the excitation and emission spectra of each probe fluorescence could be specifically collected without bleed-through. All sample mounting conditions and image acquisition parameters at 20x (NA 0.75) or 40x oil (NA 1.30) were standardized to allow better signal intensity comparison of the different tissue samples. Magnifications or zoomed in regions of interest were all imaged at similar pixel density (512×512). The images were collected as single z-planes that correspond to 2.0 µm in z depth for 20x magnification and 1.0 µm in z depth for 40x magnification.

## Results

### Virus prevalence and levels across the different developmental stages of the medfly

Normalized viral RNA levels were assessed for 13 medfly RNA viruses in larvae, pupae, and adults of the laboratory-reared Madrid strain and the mass-reared V8A strain. Three positive ssRNA viruses were detected in both medfly strains: Ceratitis capitata iflavirus 2 (CcaIV2), Ceratitis capitata iflavirus 4 (CcaIV4), and Ceratitis capitata nora virus (CcaNV). Additionally, Ceratitis capitata narnavirus (CcaNaV), and Ceratitis capitata nodavirus (CcaNdV) were detected in the Madrid strain (Figure 1A). Most viruses presented similar viral RNA levels in the different developmental stages (Figure 1). Instead, CcaNdV levels were significantly higher in the adults of the Madrid strain respective to the pupae (P = 0.0350) and the larvae (P = 0.0041) (Figure 1A, Table S3). Similarly, CcaNV levels were significantly higher in the adults of the V8A strain than the larvae (P = 0.0175), although CcaNV levels greatly varied between the adult samples (Figure 1B, Table S3).

**Figure 1.**
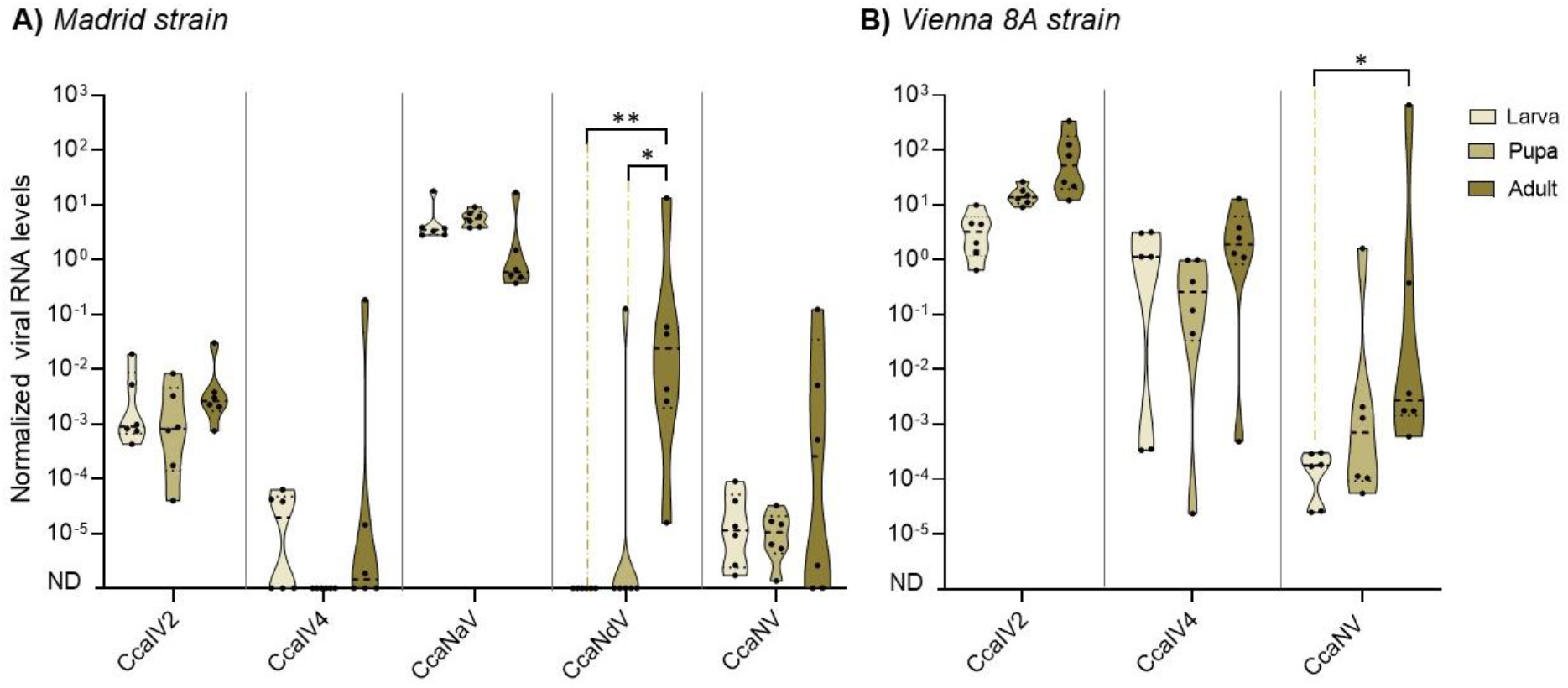
Comparison of viral RNA levels across medfly developmental stages. Normalized viral RNA levels were assessed in medfly larvae, pupae, and adults of the Madrid strain (A) and the V8A strain (B). Six biological samples each representing pools of five individuals were analyzed per developmental stage for each medfly strain. Normalized viral RNA levels were calculated by comparing the RNA levels of each virus with the transcript levels of the medfly *L23a* ribosomal gene. The absence of a virus in a specific sample was represented as non-detected (ND) in the figure. Only viruses that were detected in at least one of the samples are shown. Statistical differences between conditions are represented using asterisks (* = P < 0.05, ** = P < 0.01).

Additionally, viral levels were individually assessed in 18 adults of the Madrid and V8A strains to get insight into the prevalence of RNA viruses in adult samples (Figure 2). In the Madrid strain CcaNaV was present in most of the individuals (17 out of 18) and presented the highest viral RNA levels (Figure 2A) as observed in the analysis of developmental stages (Figure 1). CcaIV2 and CcaNV were also detected at high prevalence (17 and 15 out of 18 samples, respectively), although they presented lower viral RNA levels. CcaIV4 and CcaNdV were detected in a few samples and showed viral RNA levels in the range of CcaIV2 and CcaNV (Figure 2A). In the V8A strain, both CcaIV2 and CcaIV4 displayed the highest viral RNA levels and prevalence (18 and 16 out of 18 samples, respectively; Figure 2B). CcaNV was detected in 9 out 18 individuals and showed lower viral RNA levels. In contrast to the analysis of developmental stages, in which CcaNaV and CcaNdV were not detected in the V8A strain (Figure 1), low levels of CcaNaV and CcaNdV were detected in five and one V8A individual adults, respectively (Figure 2B).

**Figure 2.**
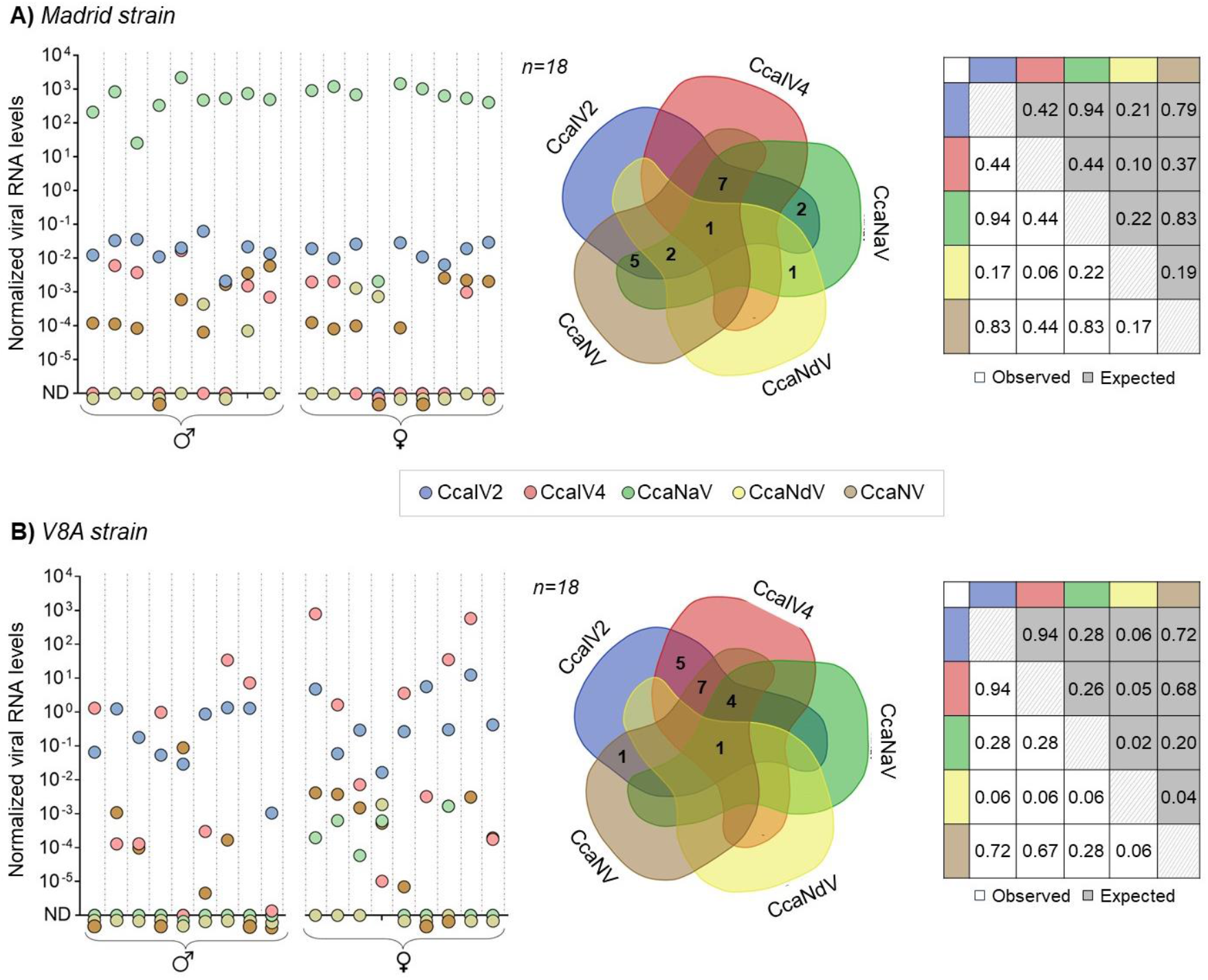
Viral prevalence and levels of RNA viruses in medfly adults. Nine adult males and nine adult females were analysed per strain: Madrid (A) and V8A (B). The dot graphs show the normalized viral RNA levels calculated by comparing the levels of each virus with the levels of the medfly *L23a* ribosomal gene. The absence of a virus in a specific sample was represented as non-detected (ND) in the figure. Venn Diagrams show all the possible combinations of RNA viruses detected in the above-mentioned samples. The coloured tables compare the expected and observed proportions of each viral combination.

Viral prevalence results were further analysed to unravel whether some viral pairwise combinations were favoured or disfavoured in the medflies. All five viruses were found to co-occur in only two individuals, one per strain. Instead, no single infected individuals were identified in any strain (Figure 2). In the Madrid strain, the most common co-occurrence was of CcaIV2, CcaIV4, CcaNaV, CcaNV (found together in seven individuals), and most individuals were co-infected by three or more viruses (15/18 individuals). In the V8A strain, the most common co-occurrence was of CcaIV2, CcaIV4, CcaNV (found together in seven individuals), and six individuals were infected with only two viruses, five of them with a combination of CcaIV2 and CcaIV4. In addition, the observed proportions of pairwise viral combinations (proportion of individuals sharing a pair of viruses in a population) matched the expected proportions (result of multiplying the individual proportions of each virus in a population) in both Madrid and V8A medfly strains (Fishe’s exact test, P > 0.999) (Figure 2, Table S2). This may indicate that an infection with any of the found RNA viruses does not positively or negatively influence the presence and abundance of the other observed RNA viruses.

### Vertical transmission is possible for the five RNA viruses

After characterizing the repertoire of RNA viruses, we explored the possible routes of transmission determining the viral distribution observed in the Madrid and V8A medfly strains. The five viruses were identified in the eggs, confirming their vertical transmission. However, the analysis of viral RNA levels in sterilized and non-sterilized eggs revealed two different transmission patterns. On the one hand, the high levels of CcaIV2, CcaIV4 and CcaNaV found for both sterilized and non-sterilized medfly eggs indicated transovarial vertical transmission of these viruses (Figure 3, Table S2). On the other hand, CcaNdV and CcaNV levels were significantly higher in non-sterilized eggs for both medfly strains, suggesting that transovum vertical transmission (associated to the eggshell) is most likely for these viruses (Figure 3, Table S2). However, the presence of low levels of CcaNdV and CcaNV in sterilized eggs of the Madrid strain may indicate that transovarial vertical transmission is also possible (Figure 3).

**Figure 3.**
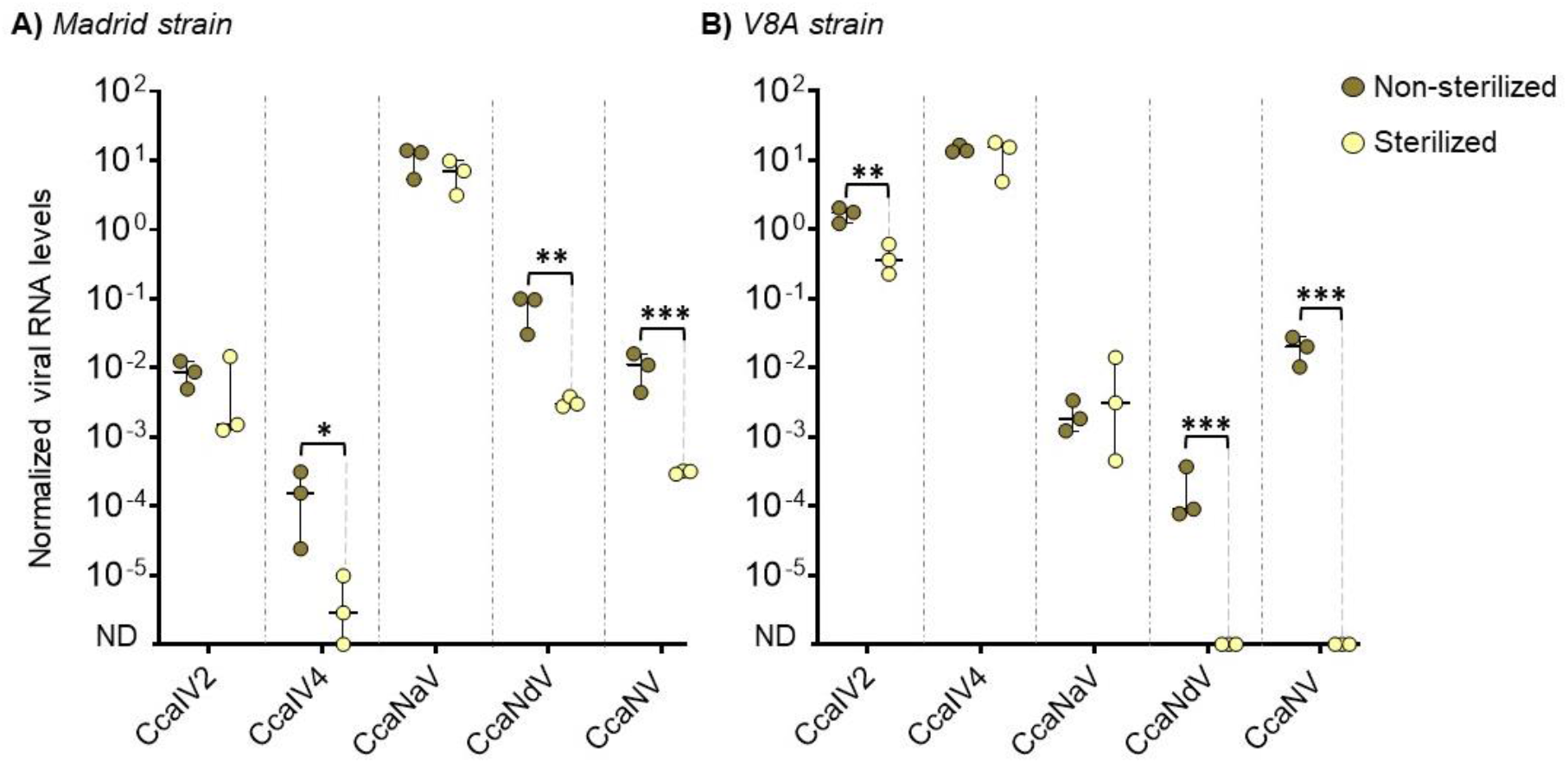
Analysis of vertical transmission via eggs. Normalized viral RNA levels in sterilized and non-sterilized eggs from the Madrid strain (A) and the V8A strain (B). Three biological samples representing pools of 0.1 ml of eggs were analyzed per strain. Normalized viral RNA levels were calculated by comparing the RNA levels of each virus with the transcript levels of the medfly *L23a* ribosomal gene as endogenous control. The absence of a virus in a specific sample was represented as non-detected (ND) in the figure. Statistical differences between conditions are represented using asterisks (* = P < 0.05, ** = P < 0.01, *** = P < 0.001).

Additionally, fifteen mating pairs consisting of a V8A male and a Madrid female, and fifteen reciprocal mating pairs consisting of a Madrid male and a V8A female were established. Of the thirty mating pairs, twenty-two produced eggs, eleven per combination. Viral RNA levels were determined in the adults forming the successful mating pairs (Figure S3 and Figure 4), and five mating pairs per group were selected to analyze the viral RNA levels in six individuals of their offspring (Figure 4). As previously observed in the analysis of viral transmission via the eggs, the results of the mating experiments evidenced two distinct transmission patterns.

**Figure 4.**
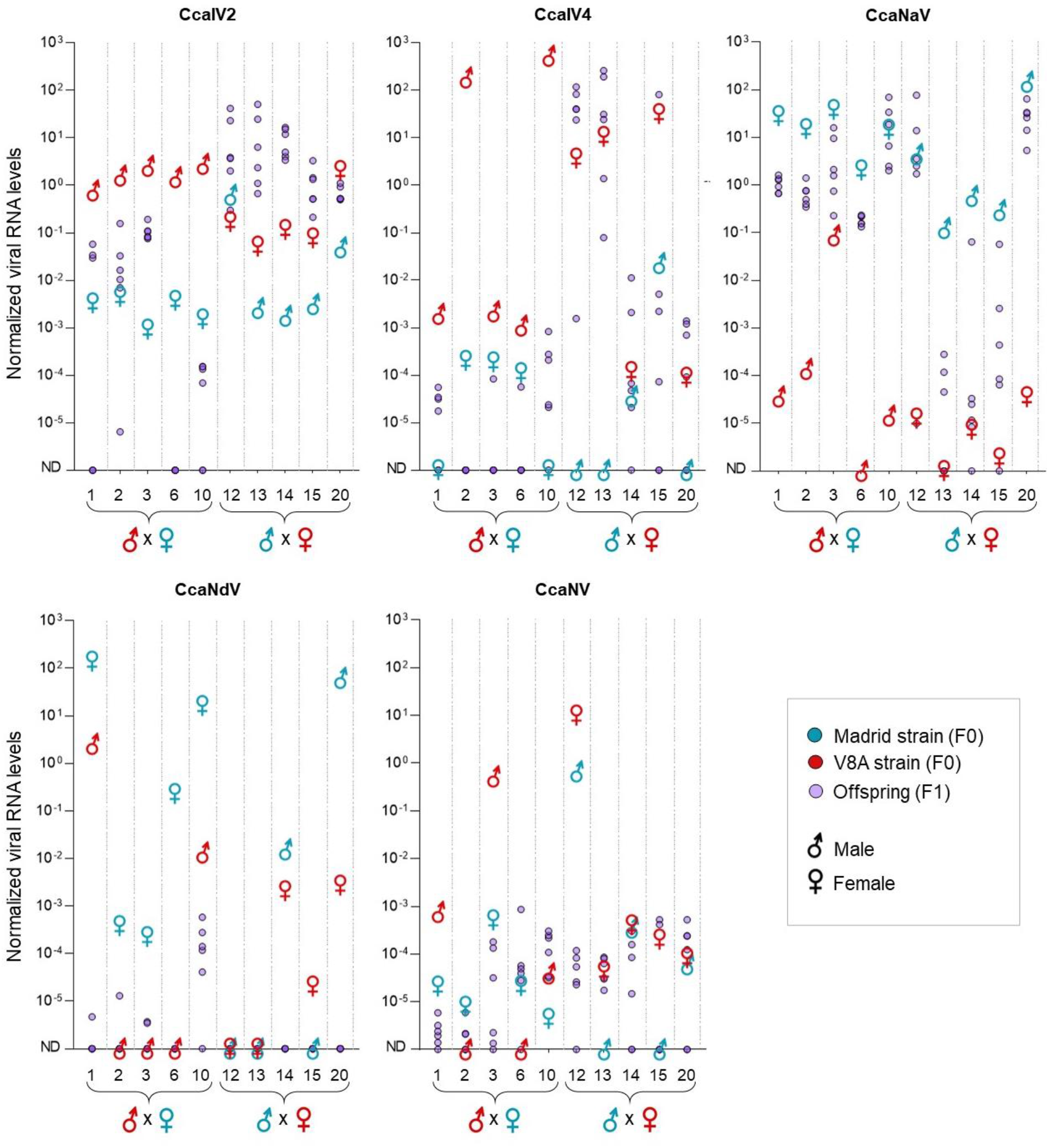
Transmission of RNA viruses to the offspring of single mating pairs. Normalized viral RNA levels in males and females forming the mating pairs and in six individuals of their offspring. Out of the ten mating pairs analyzed, half consisted of a Madrid female with a V8A male, and vice versa, as indicated at the bottom of the graph. Five graphs are shown, one per virus. Normalized viral RNA levels were calculated by comparing the RNA levels of each virus with the transcript levels of the medfly *L23a* ribosomal gene as endogenous control. The absence of a virus in a specific sample was represented as non-detected (ND) in the figure.

For CcaIV2, CcaIV4 and CcaNaV the viral presence and levels in the offspring is in most cases linked to the viral RNA levels in the female parent. CcaIV2 was detected in all the adults forming the mating pairs although at higher viral RNA levels in the V8A strain (Figure S3). A complete CcaIV2 transmission was observed in the offspring of V8A females. Instead, CcaIV2 was only detected in 18/30 descendants of Madrid females, with mating pair number 6 fully failing to transmit CcaIV2 to the offspring (0/6) (Figure 4). Similarly, CcaIV4 was successfully transmitted to the offspring (24/30) of CcaIV4 infected V8A females (Figure 4). Instead, transmission efficiency was lower for the mating pairs formed by CcaIV4-infected V8A males and Madrid females (11/30). For instance, the male parent of pair number 2 showed high levels of CcaIV4 but failed to transmit CcaIV4 to the offspring. However, the presence of CcaIV4 in the offspring of mating pairs number 1 and number 10, in which the virus was absent in the female parent, confirmed that CcaIV4 can be vertically transmitted via the male (Figure 4). For CcaNaV, which was ubiquitous in the adults of the Madrid strain (Figure S3), the vertical transmission was complete (30/30) for the offspring of CcaNaV-infected females, and incomplete (25/30) for the offspring of CcaNaV-infected males. As for CcaIV4, the presence of CcaNaV in the offspring of pairs number 13, 14, and 15, in which CcaNaV was absent or at very low levels in the female parent, indicated that vertical transmission via the male is possible (Figure 4).

On the other side, our results indicated a relatively low efficiency of vertical transmission for CcaNdV and CcaNV compared to the iflaviruses and the narnavirus. CcaNdV was only detected in 9/60 individuals of the offspring, five of them deriving from mating pair number 10 (Figure 4), despite its prevalence in the adults of the Madrid strain being higher than 50% (Figure S3). For instance, the high CcaNdV levels detected in females of mating pairs number 1 and 6 resulted in no or very low CcaNdV levels in the offspring. Meanwhile, CcaNV was identified in both Madrid and V8A adults forming the mating pairs, with higher viral RNA levels in the V8A strain for most of the mating pairs and with a high variability between samples (Figure 4, S3). After mating, CcaNV was detected in 45/60 individuals of the offspring, although at relatively low levels. Specifically, descendants of pair number 12 showed low CcaNV levels despite the high CcaNV levels detected in their parents (Figure 4).

Overall, our results suggest that vertical transmission is the main route employed by iflaviruses and narnavirus, with a preference for vertical transmission via the female. Instead, the inefficient vertical transmission observed for CcaNdV and CcaNV suggests that horizontal transmission may be the key route for nodavirus and nora virus transmission and maintenance in medfly populations.

### Horizontal transmission is likely for CcaNV and CcaNdV

During crossing experiments, co-habitation and mating may lead to the transmission of viral particles between the individuals forming the mating pairs. To test this hypothesis, we selected four mating pairs per virus in which horizontal transmission was suspected based on the viral RNA levels found in the adults. The presence of the viral negative strand was tested as an indicator of viral replication in the infected adult and the suspected recipient adult. In this assay, those suspected recipient adults in which the negative strand was detected were counted as being infected due to horizontal transmission during the nine days of co-habitation. Instead, those suspected recipient adults in which no negative strand was detected were counted as not successfully infected, pointing out to surface contamination.

For CcaIV2, the viral negative strand was detected in both parents of the samples under analysis, confirming that CcaIV2 is actively replicating in both Madrid and V8A strains (Figure 5). In this context, the presence of CcaIV2 in both medfly strains complicated the drawing of conclusions about horizontal transmission of this virus. The negative strand of CcaIV4 was detected in the V8A flies under study and in none but one sample of the Madrid strain, indicating that horizontal transmission of CcaIV4 is likely here but limited (Figure 5). However, based on previous results on CcaIV4 prevalence in the Madrid strain (Figure 2, Figure 4), we cannot exclude that the Madrid strain male of pair number 15 was possibly infected with CcaIV4 before co-habitation. The negative strand of CcaNaV was only amplified in Madrid flies (Figure 5), even though CcaNaV was detected through qPCR in three V8A adults (Figure 4). This suggests that the virus was horizontally transmitted during mating but did not establish in the recipient adult (i.e., it is a temporary contamination with the viral particles after mating).

**Figure 5.**
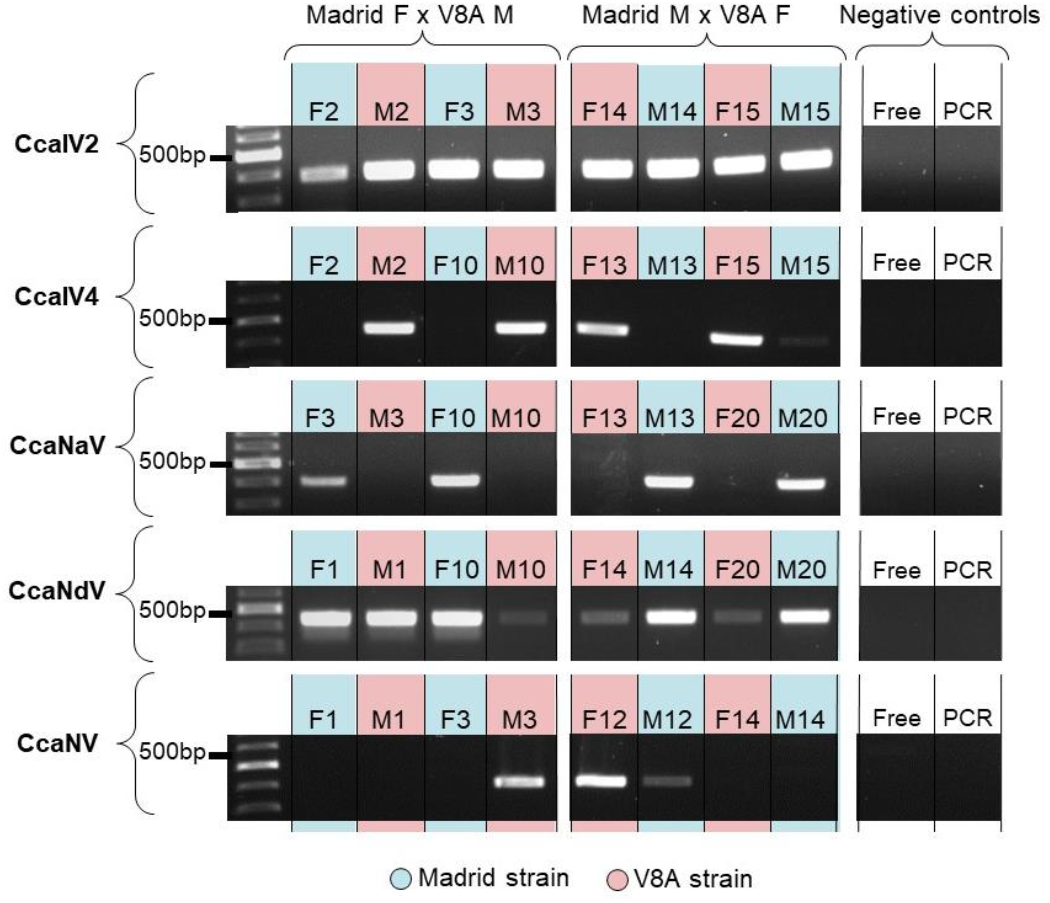
Replication of RNA viruses during co-habitation experiments. PCR amplification of the negative strand of five RNA viruses, as an indicator of viral replication. Four mating pairs formed by Madrid females (F) and V8A males (M) or *vice versa* were selected for the analysis. Two negative controls were added, one with RNA of a sample free of the virus (Free) and one without RNA (PCR). Additional negative controls can be found in Supplementary Figure 3. PCR products between 400-500 bp indicate the presence of the viral negative strand, as a result of the RT-PCR reaction using viral RNA and the primers described in Supplementary Table S1.

For CcaNdV, the negative strand was amplified in all individuals of the Madrid strain, in accordance with previous results showing higher CcaNdV levels in this strain (Figure 5, Figure 4). Instead, the presence of CcaNdV negative strand and the high viral RNA levels observed in both adults of pairs number 1, 10, 14, and 20 differed with the previous observation of low prevalence and levels of CcaNdV in the V8A strain (Figure 2). According to these results, it is tempting to hypothesize that the unexpected high CcaNdV levels and CcaNdV negative strand presence resulted from the successful CcaNdV venereal horizontal transmission between infected Madrid flies and uninfected V8A flies. Regarding CcaNV, the negative strand was only amplified in the V8A male of mating pair number 3 and both the male and female of pair number 12 (Figure 5). Interestingly, these three samples showed the highest CcaNV levels of all the mating pairs (Figure 4). In this context, the unusual high CcaNV levels in the Madrid male of mating pair number 12 may result from venereal horizontal transmission of CcaNV from the infected V8A female. However, the high variability observed for CcaNV levels in both populations (Figure 2) makes it difficult to draw solid conclusions.

### Virus tropism supports the occurrence of different transmission routes

To increase our understanding of viral tissue tropism and to explore whether tissue tropism could support the found routes of transmission, the presence of RNA viruses in gut, crop, legs, brain, ovaries, and testes of the medflies was determined by RT-qPCR.

Iflavirus and narnavirus infections were systemic in the flies, being detected in all analyzed tissues, although viral RNA levels tended to be lower in sexual tissues. For instance, CcaIV2 was less abundant in the ovaries of the V8A strain than in the legs (P = 0.0003), the brain (P = 0.0014), and the gut (P = 0.0159); and less abundant in the testes than in the legs (P = 0.0081), and the brain (P = 0.0298). Similarly, CcaIV4 was less abundant in the ovaries of the V8A strain than in the legs (P = 0.0244), the brain (P = 0.0124), and the testes (P = 0.0075) (Figure 6B, Table S4). Moreover, CcaIV4 was absent in the ovaries, the testes, and the brain of the Madrid strain, in concordance with the low CcaIV4 levels previously assessed by RT-qPCR in this strain. Regarding CcaNaV, higher normalized viral RNA levels were only identified in the brain when compared to the gut (P = 0.0019), the ovaries (P =0.0062), and the testes (P = 0.0177) (Figure 6A, Table S4).

**Figure 6.**
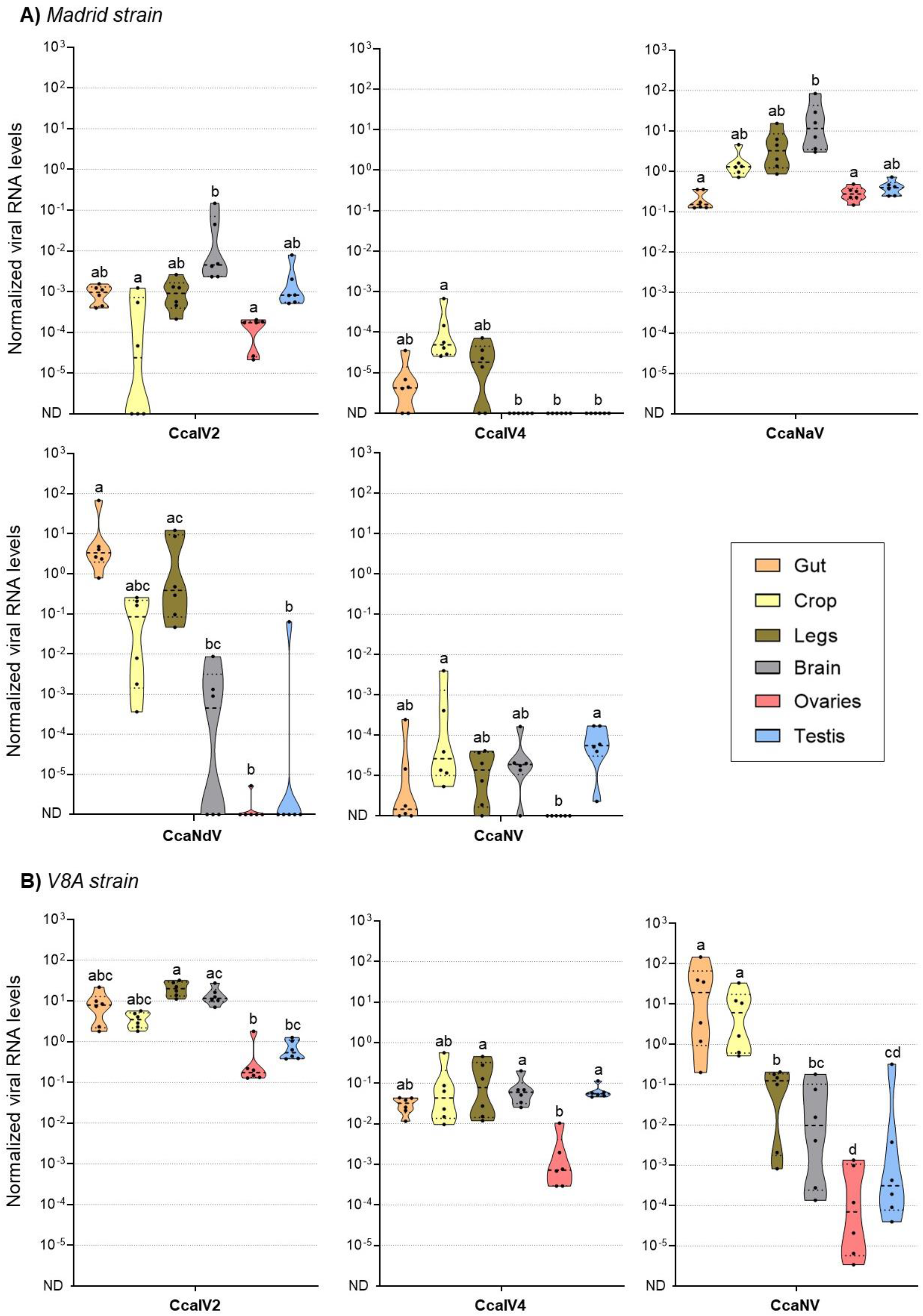
Tissue tropism of RNA viruses in adult medflies. Violin plots depict the normalized viral RNA levels across different tissues of medfly adults from the Madrid strain (A) and the V8A strain (B). Normalized viral RNA levels were calculated by comparing the RNA levels of each virus with the transcript levels of the medfly *L23a* ribosomal gene as endogenous control. The absence of a virus in a specific sample was represented as non-detected (ND) in the figure. Six samples representing pools of 8-10 identical tissues were analyzed, represented by black dots. The samples were tested for the set of viruses previously detected in the adults of each medfly strain (see Figure 1). Samples marked with the same low-case letter do not present significant differences in normalized viral RNA levels (P < 0.01).

For CcaNdV (in Madrid) and CcaNV (in V8A), higher levels were observed in the tissues related with food intake (Figure 6), supporting the preference for horizontal transmission of these viruses described above. Normalized CcaNdV levels in the Madrid strain were higher in the gut compared to the brain (P = 0.0154), the ovaries (P = 0.001), and the testes (P = 0.0025) (Table S4). In fact, CcaNdV was only detected in three out of the six samples of brain tissue, and in one sample of ovaries and testes, indicating that the infection of these tissues is possible, but not crucial for the virus (Figure 6A). Similarly, normalized CcaNV levels in the V8A strain were higher in the gut and the crop than the legs, the brain, the ovaries, and the testes (P < 0.0001) (Figure 6B, Table S4). Despite the low CcaNV levels observed in the Madrid strain, the absence of CcaNV in the ovaries generated significant differences with the crop (P = 0.0239) and the testes (P = 0.0038) (Figure 6A, Table S4).

### In situ observation of viral RNA supports the different tranmission of CcaIV2 and CcaNV

To gain insight into the localization of two RNA viruses within the medfly tissues we used the Stellaris® RNA fluorescence *in situ* hybridization (FISH) technique. From the previous results, we selected a virus that is most likely vertically transmitted (CcaIV2) and a virus that is most likely horizontally transmitted (CcaNV). CcaIV2 was also selected because of its ubiquitous distribution in diverse medfly populations (Hernández-Pelegrín et al., 2022), and CcaNV due to its association with detrimental effects during medfly development (Hernández-Pelegrín et al., 2023). Moreover, the V8A strain was selected for this analysis, since high CcaIV2 and CcaNV levels were detected in this strain by RT-qPCR (Figure 2).

Our results confirmed the differences in tissue tropism (observed with RT-qPCR) between CcaNV and CcaIV2. CcaIV2 was ubiquitously distributed in the flies, being present in all the tissues except for the crop, in which we failed to find any viral signal through FISH (Figure S4). Both male and female reproductive organs exhibited a strong CcaIV2 signal (Figure 7), supporting the hypothesis that CcaIV2 is mainly vertically transmitted. In the ovaries, CcaIV2 was localized in the ovarioles, surrounding the nucleus of the developing oocytes (Figure 7A). In the testes, CcaIV2 was mainly localized in the cells attached to the basal epithelium, where free formed spermatozoa were also observed (Figure 7B). In addition, CcaIV2 was also identified in a few cells of the vas deferens conduct, which connects the testes with the ejaculatory duct (Figure 7B.2).

**Figure 7.**
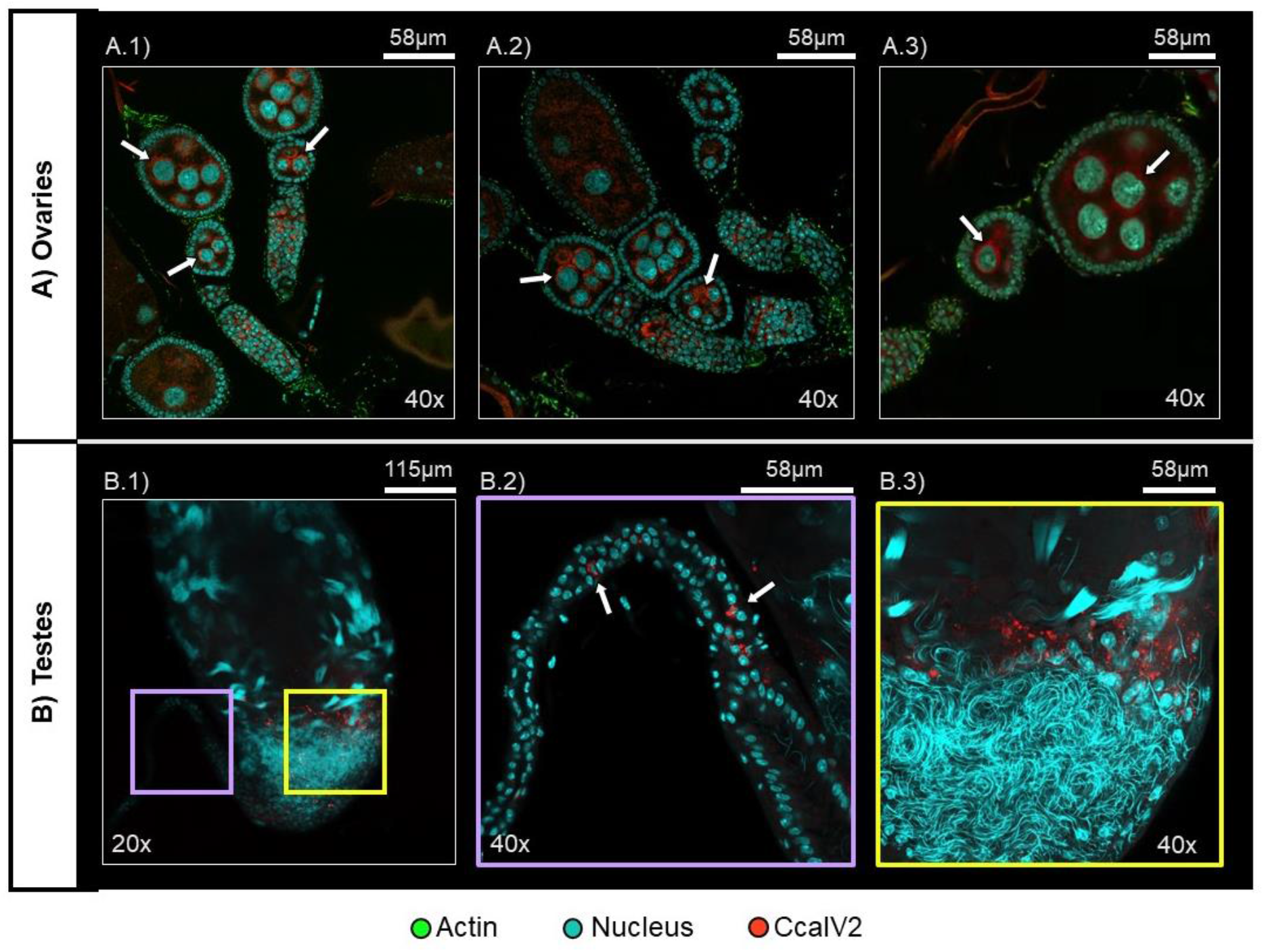
*In situ* visualization of RNA viruses in adult medfly tissues. Representative confocal planes of CcaIV2 in A) ovaries (female) and B) testes (male). Scale (in µm) and magnification (20x or 40x) are indicated for each image. B.2) and B.3) show an enlarged image of the sections of B.1) indicated with purple and yellow squares, respectively. White arrows indicate the presence of viral signal (z= 1.0 µm).

Regarding the somatic tissues, we visualized the presence of CcaIV2 infection in the gut, brain, and legs (Figure 8). In the gut, CcaIV2 density varied between samples with some guts being free of the virus and others presenting the CcaIV2 signal across the different gut segments (Figure 8A). In the brain, CcaIV2 was detected across the whole tissue, although some samples suggested that CcaIV2 levels may be higher in the central brain section (Figure 8B). In the legs, CcaIV2 was detected as a bright signal but only in a thin layer of cells close to the leg joint (Figure 8C), despite the legs presenting the highest normalized CcaIV2 levels based on RT-qPCR results (Figure 6).

**Figure 8.**
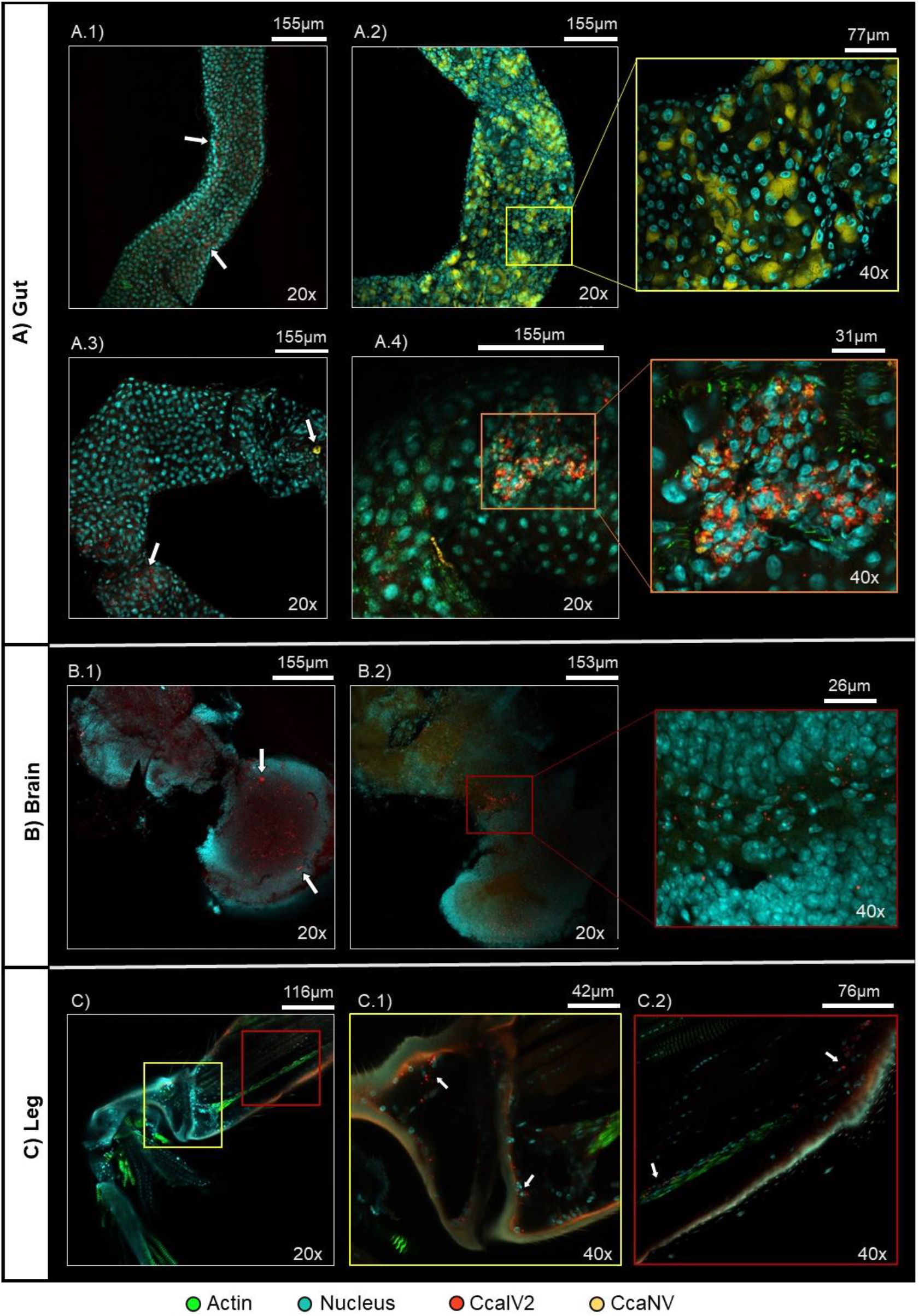
*In situ* visualization of RNA viruses in adult medfly tissues. Representative confocal images (z = 1.0 - 2.0 µm) of CcaIV2 and CcaNV visualization in three somatic tissues of the medflies: A) gut, B) brain, and C) legs. Scale (µm) and magnification (20x or 40x) are indicated for each image. Colored squares show magnified images of the sections delimited on A.2, A.4, B.2 and C, respectively.

In contrast to CcaIV2, CcaNV was predominantly detected in the gut. This agrees with the higher CcaNV levels observed in this tissue via RT-qPCR and supports the horizontal transmission of the virus. In fact, CcaNV affected most of the gut cells in some of the samples, indicated by fluorescent signal (yellow) covering the whole cytoplasm of the infected cells (Figure 8A.2). In other cases, CcaNV was absent or restricted to a few gut cells (Figure 8A.3). In contrast to our RT-qPCR results, CcaNV was not observed in the brain, legs, ovaries, and testes. This observation may be explained by the higher detection sensitivity of RT-qPCR in comparison to the FISH localization.

Since both CcaIV2 and CcaNV were detected in the gut, we screened the gut tissues for cells with CcaIV2/CcaNV co-infections. We identified one sample containing a cluster of gut cells that appeared to be co-infected with CcaIV2 and CcaNV (Figure 8A.4). In this cluster, the nuclei are smaller and aggregated, unlike the nuclei of non-infected or singly infected cells. Interestingly, apart from the cluster of cells exhibiting the suspected co-infection, very few cells of this gut were infected with any of the two viruses (Figure 8A.4).

## Discussion

Thirteen RNA viruses have been described in the medfly so far. Five of these were detected in the laboratory reared Madrid strain (CcaIV2, CcaIV4, CcaNaV, CcaNdV, CcaNV), three of which were also detected in the mass-reared V8A strain (CcaIV2, CcaIV4, CcaNV). In agreement with previous results, viral RNA levels varied between and within these medfly strains, with incomplete viral transmission patterns, except for CcaNaV in the Madrid strain and CcaIV2 in both strains (Hernández-Pelegrín et al., 2022, 2023). The ubiquitous presence of CcaIV2 is in line with previous results indicating that CcaIV2 belongs to the core RNA virome of the medfly (Hernández-Pelegrín et al., 2022). Conversely, Ceratitis capitata negev-like virus 1 was absent in all the analysed samples even though it was previously included in the medfly core RNA virome (Hernández-Pelegrín et al., 2022). This result supports the notion of a dynamic viral population in which the levels of RNA viruses can fluctuate, probably in response to environmental factors (Shapiro-Ilan et al., 2012). For instance, the direct interaction between viruses co-infecting an insect host might influence the transmission of each of those viruses. As an example, the presence of ISVs infecting mosquitoes can negatively affect the replication and transmission of arboviruses (Carvalho & Long, 2021). However, in this work we have observed that up to five viruses simultaneously co-infected the same individual and the observed ratios of pairwise viral combinations matched what was expected according to the individual prevalence of each virus. Moreover, the analysis of viral tissue tropism confirmed that different RNA viruses can co-infect the same tissues, and in some cases the same cells, as was observed for CcaIV2 and CcaNV through fluorescence *in situ* hybridization. These co-infected gut cells presented smaller and aggregated nuclei. Whether this feature results from a modification of cell division or differentiation mediated by the viruses will require further investigation. Moreover, only two RNA viruses (CcaIV2 and CcaNV) were visualized through FISH so co-infections with other RNA viruses in single cells remain unexplored.

Despite the vast variety of RNA viruses discovered in insects in the recent years, in-depth characterization of viral localization and transmission is often missing, but contributes to a better understanding of the dynamics of virus-host interactions. Our results suggest that the iflaviruses and narnavirus are mainly vertically transmitted, while the nodavirus and nora virus are preferentially horizontally transmitted. Iflaviruses are widely known to infect insects of different orders, and these infections range from covert and asymptomatic infections to infections causing clear disease symptoms (Valles et al., 2017; van Oers, 2010). Vertical and horizontal transmission routes have been characterized for iflaviruses infecting honeybees (Shen et al., 2005; Yue et al., 2006), aphids (Hatfill et al., 1990), moths (Carrillo-Tripp et al., 2014; Ponnuvel et al., 2022) and fruit flies (Morrow et al., 2023). In the medfly, the ubiquitous distribution of CcaIV2 in all developmental stages and tissues, including ovaries and testes, suggest vertical transmission of this iflavirus, as has also been reported for other iflaviruses, including Antheraea mylitta iflavirus (Ponnuvel et al., 2022) and Lymantria dispar iflavirus 1 (Carrillo-Tripp et al., 2014). Specifically, the similar CcaIV2 levels retrieved from both sterilized and non-sterilized medfly eggs strongly indicate transovarial vertical transmission, and the higher normalized CcaIV2 levels observed in the offspring of infected females implies that vertical transmission via the female is more efficient, as was also observed for an iflavirus infecting the Queensland fruit fly (Morrow et al., 2023). However, the presence of CcaIV2 in the testis, including few cells of the deferens conduct, indicate that male transmission is also possible. The natural presence of CcaIV2 in the two medfly strains made it difficult to conclude whether this virus is also horizontally transmitted. Like CcaIV2, CcaIV4 was detected in all developmental stages and tissues, although it presented higher variability and lower prevalence compared to CcaIV2. CcaIV4 was detected at similar levels in sterilized and non-sterilized eggs and was more efficiently transmitted to the offspring when mothers were infected. However, CcaIV4 was detected in the offspring of infected males and non-infected females, confirming that vertical transmission via the male is also possible. As for CcaIV2, no proof of horizontal transmission could be retrieved for CcaIV4.

The narnavirus present in the Madrid medfly strain, CcaNaV, belongs to a family of naked RNA viruses which cause asymptomatic infections in protists, fungi, plants, nematodes, or insects (Dinan et al., 2020; Espino-Vázquez et al., 2020; Hillman & Cai, 2013). The absence of extracellular virions suggests that vertical transmission is the most likely mechanism, although this has been only confirmed in *Caenorhabditis* nematodes (Richaud et al., 2019). In the medfly, CcaNaV was detected in all developmental stages and adult tissues, presented high viral RNA levels in sterilized and non-sterilized eggs, and was more efficiently transmitted by infected females than by infected males. Even though the virus was detected via RT-qPCR in initially non-infected V8A adults after mating with infected Madrid adults, the lack of detection of the negative strand indicated that the horizontal transmission did not lead to successfully established infections in V8A medflies. The detection of low CcaNaV levels in some V8A adults after mating may be explained by a) the presence of non-replicating virus as latent infection or as surface contamination, or b) the higher sensitivity of RT-qPCR with respect to the negative strand RT-PCR.

The nodavirus and nora virus present in the medfly are most likely mainly horizontally transmitted. Nodaviruses are small bi-partite viruses with a wide variety of hosts including fishes, shrimps, prawns and insects (Sahul Hameed et al., 2019). They represent a risk for the aquaculture industry, in which vertical transmission has been distinguished as the main mechanism for viral spreading (Yong et al., 2017). In the medflies, CcaNdV was unevenly distributed among developmental stages and tissues. CcaNdV was found in all adult samples but was absent in the larvae and detected in only one pupa sample, indicating that most CcaNdV infected adults failed to transmit the virus to the progeny during crossing experiments. Although these results do not suggest vertical transmission, the presence of CcaNdV in medfly eggs, with higher levels in the non-sterilized group, does not exclude transovum vertical transmission of CcaNdV. On the other hand, the highest CcaNdV levels were detected in the gut, which suggest horizontal transmission. In addition, after mating the negative strand of CcaNdV was detected in adults of the non-infected V8A population, suggesting that the virus was horizontally transmitted during co-habitation and mating leading to a successful infection of the recipient adults. The second virus discussed in this paragraph, CcaNV, belongs to the *Picornavirales* order. CcaNV is known to infect different insect species within the orders Lepidoptera (Jakubowska et al., 2014; Li et al., 2021), Hymenoptera (Remnant et al., 2017) and Diptera (Llopis-Giménez et al., 2017; Torres et al., 2012). The most representative member of this group, Drosophila melanogaster nora virus, is horizontally transmitted through the oral-fecal route (Habayeb et al., 2009). Also in our study, normalized CcaNV levels in medfly adults were higher in the tissues dealing with food intake. This result is in line with an earlier study demonstrating that CcaNV oral infection occurs after the addition of the purified virus to the larval diet (Hernández-Pelegrín et al., 2023). Moreover, the unexpectedly high CcaNV levels observed in a Madrid male after co-habitation and mating suggested the horizontal transmission of the virus via the infected V8A female, which was supported by detecting the viral negative strand in a Madrid male. On the contrary, CcaNV exhibited higher normalized viral RNA levels in non-sterilized eggs, indicating that there might be some level of vertical transmission, which is likely transovum. Our mating studies also suggested that vertical transmission of CcaNV is possible but is most likely not the major route of transmission.

Covert RNA viruses have not been related to fitness costs or benefits in the medfly, with the exception of CcaNV, which negatively affected pupal weight and adult longevity (Hernández-Pelegrín et al., 2023). However, other insect-specific RNA viruses have caused viral outbreaks and subsequent economic losses in mass-reared arthropod species, although in the natural settings these viruses have not been reported to interfere with insect health and most likely exist as covert infections. Deformed wing virus (*Iflaviridae*) compromises the development of infected honeybees and plays an important role in bee colony collapse (Martin & Brettell, 2019; Paxton et al., 2022) and the infection with nervous necrosis virus (*Nodaviridae*) resulted in high mortality in Grouper fish aquaculture (Kuo et al., 2011). The artificial rearing conditions found in insect mass-rearing industry favours an efficient transmission of RNA (and DNA) viruses and consequently may lead to the appearance of viral outbreaks under certain conditions. Decreasing insect densities, and a minimized number of shared resources may help to control viral horizontal transmission (Abd-Alla et al., 2013; Maciel-Vergara & Ros, 2017; Takacs et al., 2023). On the other side, the establishment of virus-free colonies through the screening of virus-free field populations or the artificial eradication of viruses would be a suitable solution to control viral vertical transmission.

In this article, we applied state-of-the-art techniques to uncover the tropism and transmission routes of five covert RNA viruses infecting the medfly. Overall, the viral distribution in the medfly may be explained by a combination of horizontal and vertical transmission, where the importance of each mode of transmission varies per virus. Moreover, our results reveal that the establishment and maintenance of covert infections with RNA viruses is independent from the transmission route and other factors may contribute to this phenomenon. We believe that our results contribute to the understanding of viral dynamics in this insect species and provide clues for a better control of viral epidemics in the medfly mass-rearing industry.

## Conclusions

Five RNA viruses were identified covertly infecting two strains of the Mediterranean fruit fly (medfly). By combining complementary approaches, we demonstrated that these RNA viruses vary in their mode of transmission, showing preference either for vertical transmission (Iflaviruses and narnavirus), or for horizontal transmission (nodavirus and nora virus). Consistently, the preference for vertical transmission was associated to a systemic infection in the adult tissues, while the preference for horizontal transmission with higher viral RNA levels in the digestive tissues. In this vein, the strong iflavirus signal visualized in the medfly reproductive organs using Fluorescence in Situ Hybridization, supports the notion of CcaIV2 being vertically transmitted. Similarly, the specific localization of the nora virus in gut cells, is in line with the hypothesis of horizontal transmission. Overall, our study provides important insights into the transmission and tissue tropism of RNA viruses in the medfly and reveals that the establishment of simultaneous covert infections occurs both in horizontally and vertically transmitted viruses.

## List of abbreviations

CcaIV2: Ceratitis capitata iflavirus 2
CcaIV4: Ceratitis capitata iflavirus 4
CcaNaV: Ceratitis capitata narnavirus 1
CcaNdV: Ceratitis capitata nodavirus
CcaNV: Ceratitis capitata nora virus
ISV: insect specific virus
PFA: paraformaldehyde
RT-qPCR: reverse transcription quantitative PCR
ssRNA: single strand RNA
V8A: Vienna 8A

## Data availability

All the data are available in the manuscript and the supplementary material.

## Author’s contributions

Luis Hernández-Pelegrín, Salvador Herrero and Vera I. D. Ros contributed to the study conception and design. Experimental data was collected by Luis Hernández-Pelegrín in collaboration with Hannah-Isadora Huditz for the fluorescence *in situ* hybridization; and with Pablo García-Castillo for the analysis of viral transmission. Norbert C.A. de Ruijter contributed to the visualization and interpretation of FISH results. The first draft of the manuscript was written by Luis Hernández-Pelegrín and reviewed by Salvador Herrero, Vera I. D. Ros and Monique M van Oers. All authors read and approved the final manuscript.

## Acknowledgements

We acknowledge Jaime García de Oteyza and Óscar Dembilio from TRAGSA, and Francisco Beitia from the “Instituto Valenciano de Investigaciones Agrarias” for providing medfly samples from their rearing facilities. We acknowledge Anton Strunov from the Medical University of Vienna for his advice concerning Fluorescence In Situ Hybridization.

