## Supplementary Figures and Tables for "Covert RNA viruses in medflies differ in their mode of transmission and tissue tropism"

**Supplementary material:**

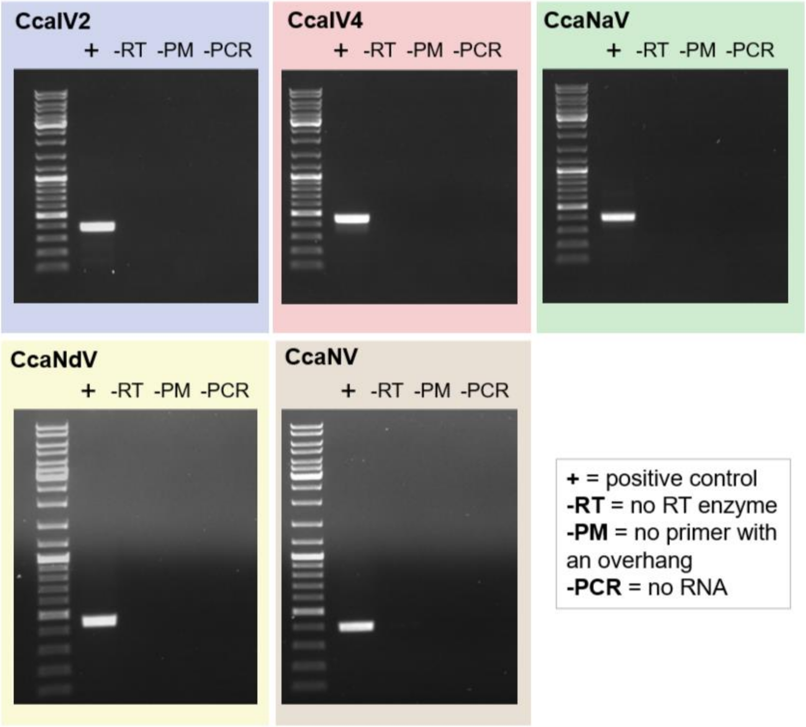


**Figure S1.** Negative controls included in the PCRs to test the accuracy of viral negative strand amplification. Three negative controls were tested per virus, involving different steps of the negative strand amplification protocol: -RT: the conversion from RNA to cDNA was performed without the RT enzyme; -TAG: the conversion from RNA to cDNA was performed without the primer with an overhang; -PCR: a PCR control in which no cDNA template is added. + represents a positive control, including a template that was known to contain the targeted virus.

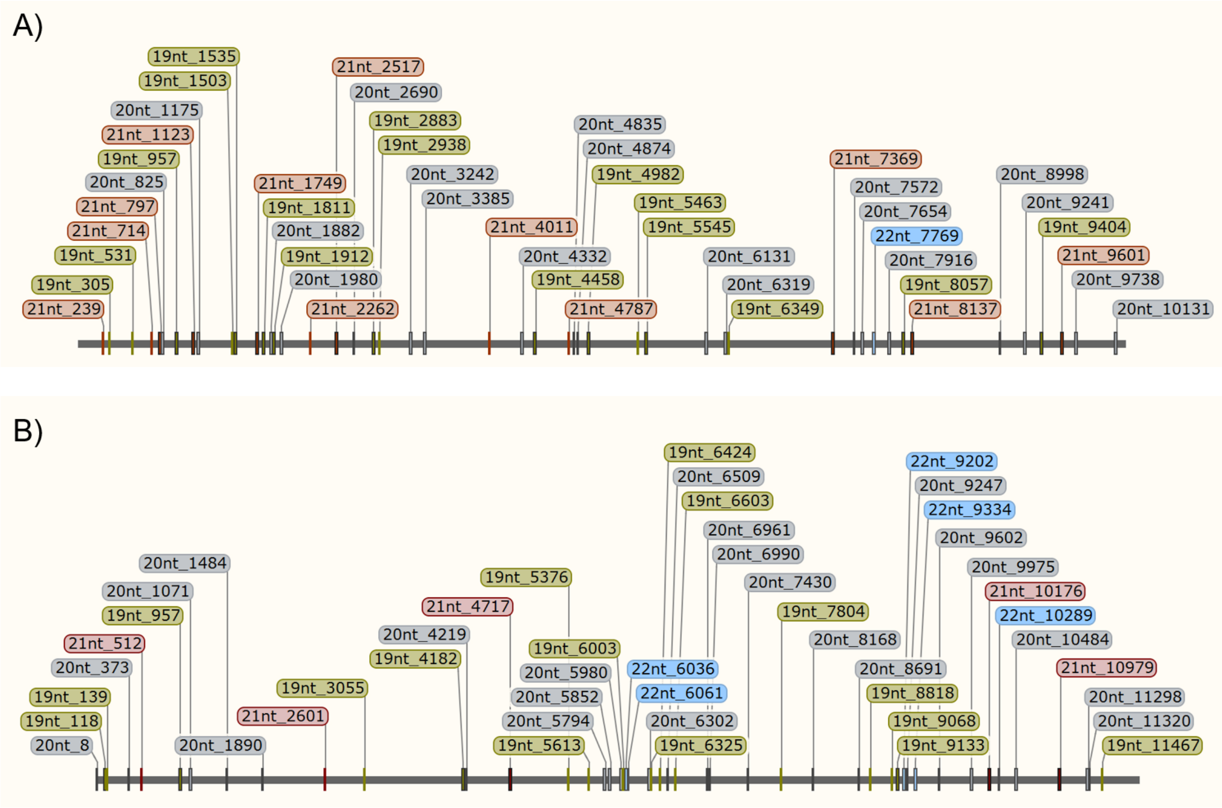


**Figure S2.** DNA probes designed for the specific visualization of Ceratitis capitata iflavirus 2 (A) and Ceratitis capitata nora virus (B) using fluorescence in situ hybridization (FISH). Probes are displayed in their mapping position along the complete genome sequences of each virus. The name of the probe contains information about its length and the starting position on the viral genome. Probes of four different lengths were designed: 19nt (dark yellow), 20nt (grey), 21nt (dark red) and 22nt (blue).

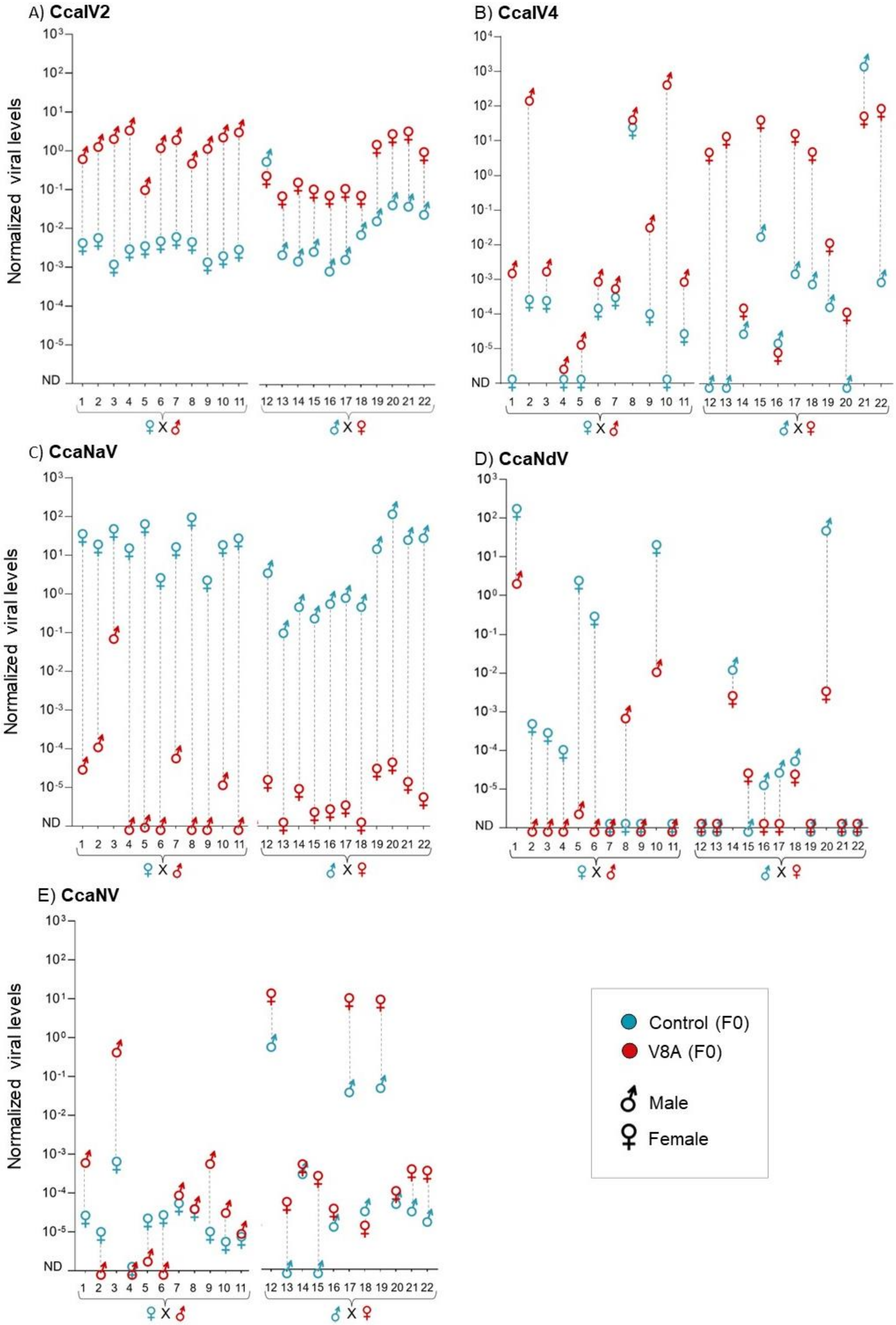
A) CcaIV2
B) CcaIV4
C) CcaNaV
D) CcaNdV
E) CcaNV
Normalized viral levels
ND
♀ X ♂
♂ X ♀
Control (F0)
V8A (F0)
Male
Female

**Figure S3.** Viral abundance in males and females forming mating pairs between the Madrid and V8A medfly strains. Out of the 22 mating pairs analyzed, half consisted of a Madrid female with a V8A male (1-11) and half of a Madrid male with a V8A female (12-22), as indicated at the bottom of the graphs. Five graphs are shown, one per virus. Normalized viral levels were calculated by comparing the levels of each virus with the levels of the medfly *L23a* ribosomal gene as endogenous control. The absence of a virus in a specific sample was represented as non-detected (ND) in the figure.

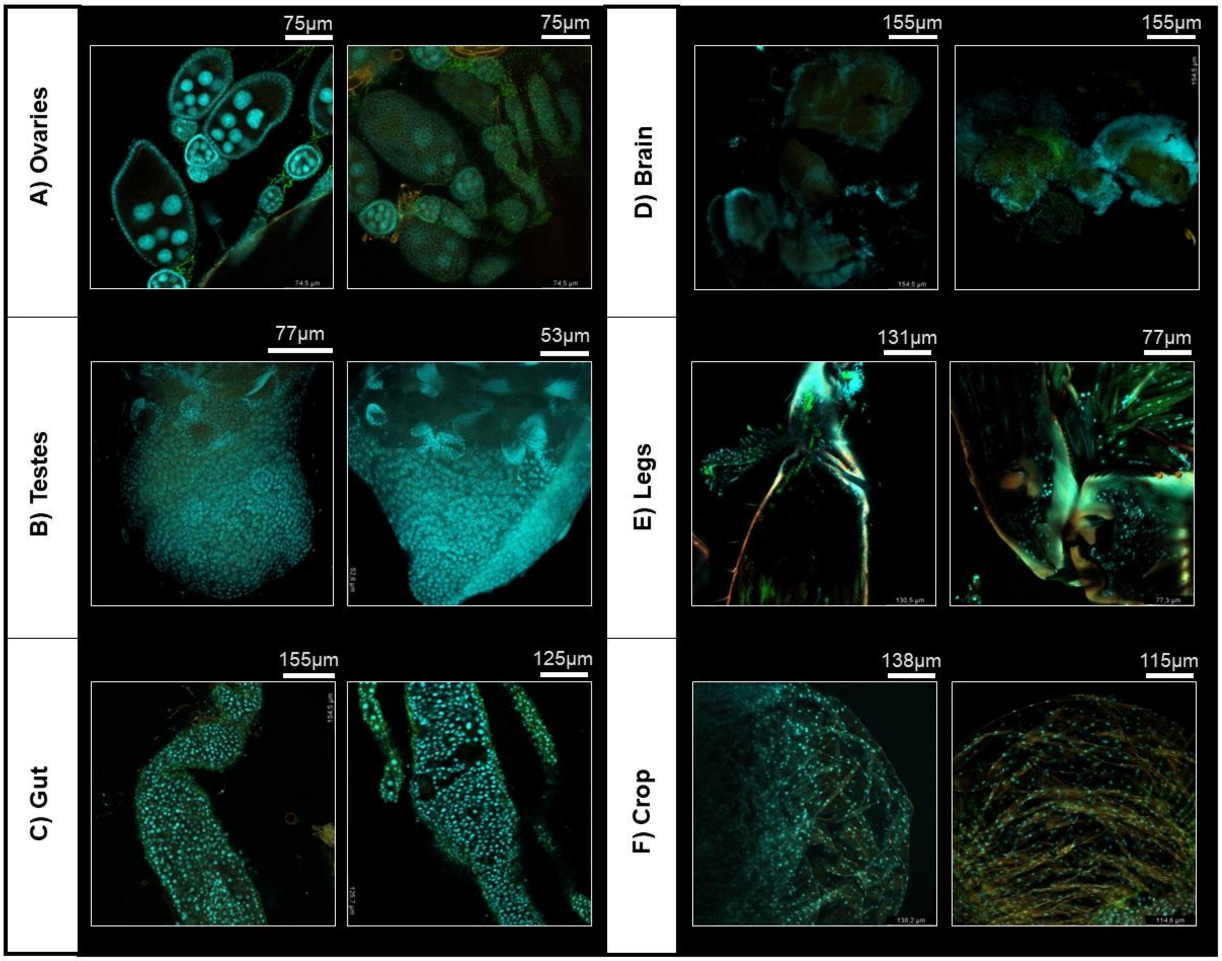


**Figure S4.** *In situ* visualization of RNA viruses in adult medfly tissues. Representative images of the visualization of medfly tissues when no CcaIV2 and CcaNV probes were added as a negative control. Scale (µm) is indicated for each image.

**Supplementary Table 1.** List of the 13 RNA viruses described in the medfly and the primers designed to amplify part of the genome of each virus through RT-qPCR (Hernández-Pelegrín et al., 2022). Primers for the reference gene L23a are also given (Llopis-Giménez et al., 2017). Primer efficiencies were calculated for the primer pairs showing viral amplification and for the reference gene. List of primers for the amplification of the negative strand of 5 RNA viruses by cDNA synthesis and PCR.

| **Virus** | **NCBI accession number** | **Forward primer (5′ to 3′)** | **Reverse primer (5′ to 3′)** | **Estimated efficiency (%)** | **R²** |
|---|---|---|---|---|---|
| **CcaIV1** | GAMC01001920.1 | TATGTGTTGTCCACCCCAGC | TGTGCGCATGGTGATTCTAAC | - | - |
| **CcaIV2** | OL957305 | CCAAGATAAGAATAGGCTAATGCGT | AACATTCGCATTTACCATTAACACAG | 93.5 | 0.993 |
| **CcaIV3** | HG994137 | GATGTTGGTGAGTTCCTC | CTATTGCCGTTAGTGAGAC | - | - |
| **CcaIV4** | HG994138 | CGTTGGTTGTTTGATTTGGAAG | GTGAAGATTGTTGGCGGAAC | 101.5 | 0.997 |
| **CcaNaV1** | OL957306 | GCTCCATCTTGGGTTCATTTG | CTGCCGTCTCCATCTTTCC | 106.1 | 0.996 |
| **CcaNdV1** | OL957308 | CTAATCGGTATGGGACAC | GACAAACTGGAGACACAAC | 105.0 | 0.945 |
| **CcaNeLV1** | HG994139 | GCGGGTAGAAGAGTATTG | GACATTCGCAGATAACAG | - | - |
| **CcaNeLV2** | OL957307 | CCCATCTAACCCTACTCC | GCGTTCTTCTTGTCATTC | - | - |
| **CcaNV** | GAMC01015827.1 | GCTCAAGCAAAAGCAGGCAG | AGCATTTCAGTCCGCTACGA | 112.4 | 0.993 |
| **CcaViLV1** | GAMC01017950.1 | ATGATACGGTTGACTTTCAG | GAGGTGGTGACGAGTAAG | - | - |
| **CcaSV** | KR822825.1 | TACAAACAAACACGAAGC | CACACCCATAACCTACAG | - | - |
| **CcaRLV1** | OL957310 | GACTAACAGGTCTCAAGG | CGCTACTCTAACTTGACC | - | - |
| **CcaTV1** | OL957313 | CTCAAGGAAGAGACACAG | GAGATATGTCTGGCGTAG | - | - |
| **L23a** | XM004518966 | GCCGAGAAATCCGCCAAATC | CTTTGCTGCAGCGGGTTTAG | 107.6 | 0.995 |

PRIMERS FOR THE AMPLIFICATION OF VIRAL NEGATIVE STRAND

| **Virus** | **NCBI accession number** | **Forward primer (5′ to 3′)** (Containing a 5′overhang in italics) | **Reverse primer (5′ to 3′)** |
|---|---|---|---|
| **CcaIV2** | OL957305 | *GGATGCAGGCTACGTGAAGATACG*CCAAGATAAGAATAGGCTAATGCGT | GGCAAGCTTTGGATTTTATCA |
| **CcaIV4** | HG994138 | *GGATGCAGGCTACGTGAAGATACG*GATGGACACAGTCGGAGCTTTAGC | AACATTCGCATTTACCATTAACACAG |
| **CcaNaV1** | OL957306 | *GGATGCAGGCTACGTGAAGATACG*CACCGCTCCATCTTGGGTTCATTTG | TGTGCGCATGGTGATTCTAAC |
| **CcaNdV1** | OL957308 | *GGATGCAGGCTACGTGAAGATACG*GCTAATCGGTATGGGACACAAGGAC | TGTTTCAACGCCGTCAGGT |
| **CcaNV** | GAMC01015827.1 | *GGATGCAGGCTACGTGAAGATACG*GCTCAAGCAAAAGCAGGCAGAACAC | CTTTGCTGCAGCGGGTTTAG |
| **Primer mapping the overhang** | | GATGCAGGCTACGTGAAGATACG | |

**Supplementary Table 2.** Statistical differences in viral levels between medfly developmental stages, between adult tissues (tissue tropism), and between sterilized versus non-sterilized eggs (vertical transmission). Test were performed per virus, in each of the two medfly strains under analysis.

| Experiment | Strain | Virus | Test | Result |
|---|---|---|---|---|
| Developmental Stages | Madrid strain | CcaIV2 | ANOVA | P = 0.2681 |
| | | CcaIV4 | Kruskal-Wallis (non-parametric) | P = 0.1763 |
| | | CcaNaV | ANOVA | P = 0.0177 |
| | | CcaNdV | Kruskal-Wallis (non-parametric) | P = 0.001 |
| | | CcaNV | Kruskal-Wallis (non-parametric) | P = 0.3485 |
| | V8A strain | CcaIV2 | ANOVA | P = 0.0002 |
| | | CcaIV4 | Kruskal-Wallis (non-parametric) | P = 0.0916 |
| | | CcaNV | Kruskal-Wallis (non-parametric) | P = 0.0143 |
| Tissue Tropism | Madrid strain | CcaIV2 | Kruskal-Wallis (non-parametric) | P = 0.0002 |
| | | CcaIV4 | Kruskal-Wallis (non-parametric) | P = 0.0001 |
| | | CcaNaV | ANOVA | P < 0.0001 |
| | | CcaNdV | Kruskal-Wallis (non-parametric) | P < 0.0001 |
| | | CcaNV | Kruskal-Wallis (non-parametric) | P = 0.0048 |
| | V8A strain | CcaIV2 | ANOVA | P < 0.0001 |
| | | CcaIV4 | Kruskal-Wallis (non-parametric) | P = 0.0037 |
| | | CcaNV | ANOVA | P < 0.0001 |
| Vertical transmission via the eggs | Madrid strain | CcaIV2 | Unpaired t-test | t=1.185, df=4 P = 0.3016 |
| | | CcaIV4 | Unpaired t-test | t=3.507, df=4 P = 0.0247 |
| | | CcaNaV | Unpaired t-test | t=1.085, df=4 P = 0.339 |
| | | CcaNdV | Unpaired t-test | t=7.551, df=4 P = 0.0016 |
| | | CcaNV | Unpaired t-test | t=8.835, df=4 P = 0.0009 |
| | V8A strain | CcaIV2 | Unpaired t-test | t=4.616, df=4 P = 0.0099 |
| | | CcaIV4 | Unpaired t-test | t=0.6440, df=4 P = 0.5547 |
| | | CcaNaV | Unpaired t-test | t=0.3128, df=4 P = 0.7701 |
| | | CcaNdV | Unpaired t-test | t=9.846, df=4 P = 0.0006 |
| | | CcaNV | Unpaired t-test | t=33.74, df=4 P < 0.0001 |
| Expected versus observed values of pairwise viral combinations | Madrid and V8A strains | CcaIV2-CcaIV4 | Fisher´s exact test | P > 0.999 |
| | | CcaIV2-CcaNaV | Fisher´s exact test | P > 0.999 |
| | | CcaIV2-CcaNdV | Fisher´s exact test | P > 0.999 |
| | | CcaIV2-CcaNV | Fisher´s exact test | P > 0.999 |
| | | CcaIV4-CcaNaV | Fisher´s exact test | P > 0.999 |
| | | CcaIV4-CcaNdV | Fisher´s exact test | P > 0.999 |
| | | CcaIV4-CcaNV | Fisher´s exact test | P > 0.999 |
| | | CcaNaV-CcaNdV | Fisher´s exact test | P > 0.999 |
| | | CcaNaV-CcaNV | Fisher´s exact test | P > 0.999 |
| | | CcaNdV-CcaNV | Fisher´s exact test | P > 0.999 |

**Supplementary Table 3.** Results of Tukey´s multiple comparisons (two-way ANOVA) or Dunn´s multiple comparisons (non-parametric Kruskal-Wallis test) between medfly developmental stages. Test were performed per virus, in each of the two medfly strains under analysis.

| Developmental Stages, Madrid strain | | | | |
|---|---|---|---|---|
| CcaIV2 (Tukey´s multiple comparisons) | Mean Diff, | 95% CI | Significance | P Value |
| Larva vs. Pupa | 0.3842 | -1.232 to 2.000 | ns | 0.8373 |
| Larva vs. Adult | -0.2788 | -1.895 to 1.337 | ns | 0.9106 |
| Pupa vs. Adult | -0.663 | -2.279 to 0.9531 | ns | 0.5911 |
| CcaIV4 (Dunn´s multiple comparisons) | Mean rank diff. | Summary | P Value | |
| Larva vs. Pupa | 4,750 | ns | 0,1992 | |
| Larva vs. Adult | 0,5000 | ns | >0,9999 | |
| Pupa vs. Adult | -4,250 | ns | 0,3015 | |
| CcaNaV (Tukey´s multiple comparisons) | Mean Diff, | 95% CI | Significance | P Value |
| Larva vs. Pupa | -0.1072 | -1.723 to 1.509 | ns | 0.9862 |
| Larva vs. Adult | 0.6079 | -1.008 to 2.224 | ns | 0.6423 |
| Pupa vs. Adult | 0.7151 | -0.9010 to 2.331 | ns | 0.543 |
| CcaNdV (Dunn´s multiple comparisons) | Mean rank diff. | Summary | P Value | |
| Larva vs. Pupa | -1,833 | ns | >0,9999 | |
| Larva vs. Adult | -8,667 | ** | 0,0041 | |
| Pupa vs. Adult | -6,833 | * | 0,0350 | |
| CcaNV (Dunn´s multiple comparisons) | Mean rank diff. | Summary | P Value | |
| Larva vs. Pupa | 0,6667 | ns | >0,9999 | |
| Larva vs. Adult | -0,1667 | ns | >0,9999 | |
| Pupa vs. Adult | -0,8333 | ns | >0,9999 | |
| Developmental Stages, V8A strain | | | | |
| CcaIV2 (Tukey´s multiple comparisons) | Mean Diff, | 95% CI | Significance | P Value |
| Larva vs. Pupa | -0.7339 | -2.724 to 1.256 | ns | 0.647 |
| Larva vs. Adult | -1.3 | -3.290 to 0.6901 | ns | 0.2632 |
| Pupa vs. Adult | -0.5663 | -2.557 to 1.424 | ns | 0.7707 |
| CcaIV4 (Dunn´s multiple comparisons) | Mean rank diff. | Summary | P Value | |
| Larva vs. Pupa | 3,833 | ns | 0,6408 | |
| Larva vs. Adult | -2,833 | ns | >0,9999 | |
| Pupa vs. Adult | -6,667 | ns | 0,0916 | |
| CcaNV (Dunn´s multiple comparisons) | Mean rank diff. | Summary | P Value | |
| Larva vs. Pupa | -3,500 | ns | 0,7684 | |
| Larva vs. Adult | -8,500 | * | 0,0175 | |
| Pupa vs. Adult | -5,000 | ns | 0,3143 | |

ns = non significant; * = P = 0.01; **** = P < 0.0001

**Supplementary Table 4.** Results of Tukey´s multiple comparisons (two-way ANOVA) or Dunn´s multiple comparisons (non-parametric Kruskal-Wallis test) between medfly adult tissues. Test were performed per virus, in each of the two medfly strains under analysis.

| Adult tissues, Madrid strain | | | | |
|---|---|---|---|---|
| CcaIV2 (Dunn´s multiple comparisons) | Mean rank diff. | Summary | P Value | |
| Gut vs. Crop | 10,83 | ns | >0,9999 | |
| Gut vs. Legs | -1,333 | ns | >0,9999 | |
| Gut vs. Brain | -13,00 | ns | 0,4881 | |
| Gut vs. Ovaries | 12,33 | ns | 0,6382 | |
| Gut vs. Testes | -2,833 | ns | >0,9999 | |
| Crop vs. Legs | -12,17 | ns | 0,6814 | |
| Crop vs. Brain | -23,83 | ** | 0,0013 | |
| Crop vs. Ovaries | 1,500 | ns | >0,9999 | |
| Crop vs. Testes | -13,67 | ns | 0,3693 | |
| Legs vs. Brain | -11,67 | ns | 0,8257 | |
| Legs vs. Ovaries | 13,67 | ns | 0,3693 | |
| Legs vs. Testes | -1,500 | ns | >0,9999 | |
| Brain vs. Ovaries | 25,33 | *** | 0,0005 | |
| Brain vs. Testes | 10,17 | ns | >0,9999 | |
| Ovaries vs. Testes | -15,17 | ns | 0,1895 | |
| CcaIV4 (Dunn´s multiple comparisons) | Mean rank diff. | Summary | P Value | |
| Gut vs. Crop | -11,33 | ns | 0,5095 | |
| Gut vs. Legs | -2,667 | ns | >0,9999 | |
| Gut vs. Brain | 9,333 | ns | >0,9999 | |
| Gut vs. Ovaries | 9,333 | ns | >0,9999 | |
| Gut vs. Testes | 9,333 | ns | >0,9999 | |
| Crop vs. Legs | 8,667 | ns | >0,9999 | |
| Crop vs. Brain | 20,67 | ** | 0,0017 | |
| Crop vs. Ovaries | 20,67 | ** | 0,0017 | |
| Crop vs. Testes | 20,67 | ** | 0,0017 | |
| Legs vs. Brain | 12,00 | ns | 0,3713 | |
| Legs vs. Ovaries | 12,00 | ns | 0,3713 | |
| Legs vs. Testes | 12,00 | ns | 0,3713 | |
| Brain vs. Ovaries | 0,000 | ns | >0,9999 | |
| Brain vs. Testes | 0,000 | ns | >0,9999 | |
| Ovaries vs. Testes | 0,000 | ns | >0,9999 | |
| CcaNaV (Tukey´s multiple comparisons) | Mean Diff, | 95% CI | Significance | P Value |
| Gut vs. Crop | -0.878 | -2.209 to 0.4532 | ns | 0.4037 |
| Gut vs. Legs | -1.221 | -2.553 to 0.1097 | ns | 0.0921 |
| Gut vs. Brain | -1.804 | -3.135 to -0.4727 | ** | 0.0019 |
| Gut vs. Ovaries | -0.156 | -1.487 to 1.175 | ns | 0.9994 |
| Gut vs. Testes | -0.3064 | -1.638 to 1.025 | ns | 0.9855 |

| | | | | |
|---|---|---|---|---|
| Crop vs. Legs | -0.3435 | -1.675 to 0.9876 | ns | 0.9759 |
| Crop vs. Brain | -0.9258 | -2.257 to 0.4053 | ns | 0.3428 |
| Crop vs. Ovaries | 0.722 | -0.6091 to 2.053 | ns | 0.6222 |
| Crop vs. Testes | 0.5716 | -0.7595 to 1.903 | ns | 0.8166 |
| Legs vs. Brain | -0.5823 | -1.913 to 0.7488 | ns | 0.8046 |
| Legs vs. Ovaries | 1.065 | -0.2656 to 2.397 | ns | 0.1962 |
| Legs vs. Testes | 0.9151 | -0.4160 to 2.246 | ns | 0.356 |
| Brain vs. Ovaries | 1.648 | 0.3167 to 2.979 | ** | 0.0062 |
| Brain vs. Testes | 1.497 | 0.1663 to 2.829 | * | 0.0177 |
| Ovaries vs. Testes | -0.1504 | -1.482 to 1.181 | ns | 0.9995 |
| CcaNdV (Dunn´s multiple comparisons) | Mean rank diff. | Summary | P Value | |
| Gut vs. Crop | 10,67 | ns | >0,9999 | |
| Gut vs. Legs | 3,833 | ns | >0,9999 | |
| Gut vs. Brain | 19,50 | * | 0,0154 | |
| Gut vs. Ovaries | 23,67 | ** | 0,0010 | |
| Gut vs. Testes | 22,33 | ** | 0,0025 | |
| Crop vs. Legs | -6,833 | ns | >0,9999 | |
| Crop vs. Brain | 8,833 | ns | >0,9999 | |
| Crop vs. Ovaries | 13,00 | ns | 0,4289 | |
| Crop vs. Testes | 11,67 | ns | 0,7420 | |
| Legs vs. Brain | 15,67 | ns | 0,1251 | |
| Legs vs. Ovaries | 19,83 | * | 0,0126 | |
| Legs vs. Testes | 18,50 | * | 0,0276 | |
| Brain vs. Ovaries | 4,167 | ns | >0,9999 | |
| Brain vs. Testes | 2,833 | ns | >0,9999 | |
| Ovaries vs. Testes | -1,333 | ns | >0,9999 | |
| CcaNV (Dunn´s multiple comparisons) | Mean rank diff. | Summary | P Value | |
| Gut vs. Crop | -9,833 | ns | >0,9999 | |
| Gut vs. Legs | -3,583 | ns | >0,9999 | |
| Gut vs. Brain | -5,917 | ns | >0,9999 | |
| Gut vs. Ovaries | 9,167 | ns | >0,9999 | |
| Gut vs. Testes | -12,83 | ns | 0,4944 | |
| Crop vs. Legs | 6,250 | ns | >0,9999 | |
| Crop vs. Brain | 3,917 | ns | >0,9999 | |
| Crop vs. Ovaries | 19,00 | * | 0,0239 | |
| Crop vs. Testes | -3,000 | ns | >0,9999 | |
| Legs vs. Brain | -2,333 | ns | >0,9999 | |
| Legs vs. Ovaries | 12,75 | ns | 0,5117 | |
| Legs vs. Testes | -9,250 | ns | >0,9999 | |
| Brain vs. Ovaries | 15,08 | ns | 0,1829 | |
| Brain vs. Testes | -6,917 | ns | >0,9999 | |
| Ovaries vs. Testes | -22,00 | ** | 0,0038 | |

| Adult tissues, V8A strain | | | | |
|---|---|---|---|---|
| CcaIV2 (Tukey´s multiple comparisons) | Mean diff. | 95.00% CI | Summary | P Value |
| Gut vs. Crop | 0.2689 | -1.204 to 1.742 | ns | 0.9882 |
| Gut vs. Legs | -0.499 | -1.972 to 0.9743 | ns | 0.847 |
| Gut vs. Brain | -0.3136 | -1.787 to 1.160 | ns | 0.9764 |
| Gut vs. Ovaries | 1.409 | -0.06455 to 2.882 | * | 0.0159 |
| Gut vs. Testes | 1.003 | -0.4705 to 2.476 | ns | 0.1802 |
| Crop vs. Legs | -0.7679 | -2.241 to 0.7054 | ns | 0.4646 |
| Crop vs. Brain | -0.5825 | -2.056 to 0.8908 | ns | 0.7429 |
| Crop vs. Ovaries | 1.14 | -0.3334 to 2.613 | ns | 0.0879 |
| Crop vs. Testes | 0.734 | -0.7393 to 2.207 | ns | 0.5158 |
| Legs vs. Brain | 0.1854 | -1.288 to 1.659 | ns | 0.9979 |
| Legs vs. Ovaries | 1.908 | 0.4345 to 3.381 | *** | 0.0003 |
| Legs vs. Testes | 1.502 | 0.02855 to 2.975 | ** | 0.0081 |
| Brain vs. Ovaries | 1.722 | 0.2491 to 3.196 | ** | 0.0014 |
| Brain vs. Testes | 1.316 | -0.1568 to 2.790 | * | 0.0298 |
| Ovaries vs. Testes | -0.4059 | -1.879 to 1.067 | ns | 0.9302 |
| CcaIV4 (Dunn´s multiple comparisons) | Mean rank diff. | Summary | P Value | |
| Gut vs. Crop | -5,000 | ns | >0,9999 | |
| Gut vs. Legs | -7,500 | ns | >0,9999 | |
| Gut vs. Brain | -8,667 | ns | >0,9999 | |
| Gut vs. Ovaries | 11,67 | ns | 0,8267 | |
| Gut vs. Testes | -9,500 | ns | >0,9999 | |
| Crop vs. Legs | -2,500 | ns | >0,9999 | |
| Crop vs. Brain | -3,667 | ns | >0,9999 | |
| Crop vs. Ovaries | 16,67 | ns | 0,0922 | |
| Crop vs. Testes | -4,500 | ns | >0,9999 | |
| Legs vs. Brain | -1,167 | ns | >0,9999 | |
| Legs vs. Ovaries | 19,17 | * | 0,0244 | |
| Legs vs. Testes | -2,000 | ns | >0,9999 | |
| Brain vs. Ovaries | 20,33 | * | 0,0124 | |
| Brain vs. Testes | -0,8333 | ns | >0,9999 | |
| Ovaries vs. Testes | -21,17 | ** | 0,0075 | |
| CcaNV (Tukey´s multiple comparisons) | Mean diff. | 95.00% CI of diff. | Summary | P Value |
| Gut vs. Crop | 0.3117 | -1.162 to 1.785 | ns | 0.9771 |
| Gut vs. Legs | 2.376 | 0.9024 to 3.849 | **** | <0.0001 |
| Gut vs. Brain | 3.119 | 1.645 to 4.592 | **** | <0.0001 |
| Gut vs. Ovaries | 5.061 | 3.588 to 6.535 | **** | <0.0001 |
| Gut vs. Testes | 3.947 | 2.474 to 5.420 | **** | <0.0001 |
| Crop vs. Legs | 2.064 | 0.5907 to 3.537 | **** | <0.0001 |
| Crop vs. Brain | 2.807 | 1.334 to 4.280 | **** | <0.0001 |
| Crop vs. Ovaries | 4.75 | 3.276 to 6.223 | **** | <0.0001 |
| Crop vs. Testes | 3.635 | 2.162 to 5.109 | **** | <0.0001 |

| | | | | |
|---|---|---|---|---|
| Legs vs. Brain | 0.743 | -0.7303 to 2.216 | ns | 0.5021 |
| Legs vs. Ovaries | 2.686 | 1.213 to 4.159 | **** | <0.0001 |
| Legs vs. Testes | 1.571 | 0.09809 to 3.045 | ** | 0.0048 |
| Brain vs. Ovaries | 1.943 | 0.4695 to 3.416 | *** | 0.0002 |
| Brain vs. Testes | 0.8284 | -0.6449 to 2.302 | ns | 0.3776 |
| Ovaries vs. Testes | -1.114 | -2.588 to 0.3589 | ns | 0.1013 |

*ns = non significant; * = $P = 0.01$; ** = $P < 0.01$; *** = $P < 0.001$; **** = $P < 0.0001$
